# Red knots in Europe - a dead end host species or a new niche for highly pathogenic avian influenza?

**DOI:** 10.1101/2024.05.21.593879

**Authors:** Jacqueline King, Anne Pohlmann, Andreas Bange, Elisabeth Horn, Bernd Hälterlein, Angele Breithaupt, Anja Globig, Anne Günther, Angie Kelm, Christian Wiedemann, Christian Grund, Karena Haecker, Stefan Garthe, Timm Harder, Martin Beer, Philipp Schwemmer

## Abstract

The 2020/2021 epidemic in Europe of highly pathogenic avian influenza virus (HPAIV) of subtype H5 surpassed all previously recorded European outbreaks in size, genotype constellations and reassortment frequency and continued into 2022 and 2023. The causative 2.3.4.4b viral lineage proved to be highly proficient with respect to reassortment with cocirculating low pathogenic AIV and seems to establish an endemic status in northern Europe. A specific HPAIV reassortant of the subtype H5N3 was detected almost exclusively in red knots (*Calidris canutus islandica)* in December 2020. It caused systemic and rapidly fatal disease leading to a singular and self-limiting mass mortality affecting about 3.500 birds in the German Wadden Sea, roughly 1% of the entire flyway population of *islandica* red knots. Phylogenetic analyses revealed that the H5N3 reassortant very likely had formed in red knots and remained confined to this species. While mechanisms of virus circulation in potential reservoir species, dynamics of spill-over and reassortment events and the roles of environmental virus sources remain to be identified, the year-round infection pressure poses severe threats to endangered avian species, and prompts adaptation of habitat and species conservation practices.

**One-Sentence Summary:** High red knot mortality in Europe (December 2020) was associated with infection of a unique genotype of HPAIV H5N3 clade 2.3.4.4b.

## Background

Between mid and end December 2020, embedded in the most devastating and protracted highly pathogenic avian influenza (HPAI) clade 2.3.3.4b H5-epidemics until then in Germany, Europe and beyond, thousands of red knots (*Calidris canutus*) were found dead or dying along the shores or birds falling literally dead from the sky in the German Wadden Sea.

Red knots, a bird species belonging to the snipe family (*Scolopacidae*), are considered one of the very few species of *Charadrii* with evidence of high rates of previous AIV infection (1, 1). Two subspecies of red knot occur in Europe: The *canutus* subspecies breeding in Siberia with a population of app. 250,000 individuals (2010-2014) and the *islandica* subspecies breeding in Greenland and Canada with a population of 310,000 – 360,000 individuals (2013-2017; (2)). The *canutus* subspecies only visits the Wadden Sea areas of the southern North Sea en route to Western Africa during its autumn (mainly July-August and October) and spring migration (May), while a proportion of the *islandica* subspecies may spend also the entire winter season from November to February in the Wadden Sea (3–6). The Wadden Sea in central Europe is one of the most important staging sites for waders such as red knot worldwide, and according to a regular monitoring scheme about 75 % of the flyway population of red knots uses this area, i.e. a maximum of ca. 350,000 individuals (both subspecies together; (5)). Red knots use high tide roosts in large flocks in various places along the entire Wadden Sea coast (6). While the trend of the *canutus* population in the Wadden Sea is stable (but in Schleswig-Holstein continuous declining since the end of the 1990s), the long-term trend 1987/1988-2019/2020 of the *islandica* population in the Wadden Sea remains negative, mainly due to the decline in the Schleswig-Holstein part, and the short-term trend 2010/2011-2019/2020 is stable (5). In the European Red List of Birds the red knot is classified in the category “near threatened” (7). In the Wadden Sea area, red knots are exclusively molluscivore and known to primarily feed on Baltic tellins (*Limecola balthica*; (8)).

Wild birds of the orders *Anseriformes* (ducks, geese, swans) and *Charadriiformes* (gulls and allies [“*Lari*”], waders [“*Charadrii*”] and auks [“*Alcae*”]) are considered the main reservoir of avian influenza viruses (AIV). In Europe, the majority of low pathogenic avian influenza viruses (LPAIV) have been found in water birds, particularly dabbling ducks (9). Prevalences of LPAIV grossly vary in shorebirds in different regions globally; while in Europe AIV have only sporadically been found in waders, LPAIV infections appear to be more widespread in *Charadrii* birds in other regions of the world (10–12). A particular AIV hotspot in waders has been reported in North American Delaware Bay, one of the largest wintering and stop-over sites of shorebirds globally (13).

Prior to the emergence of highly pathogenic avian influenza (HPAI) Goose/Guangdong/1996 (Gs/Gd) lineage, isolation of HPAI viruses from *Charadriiformes* was reported only once in 1961, when an HPAIV virus (A/tern/South Africa/61 [H5N3]) caused the death of more than 1,000 Common Terns (*Sterna hirundo*) in South Africa (14).

After the emergence and continuous evolution of Gs/Gd HPAI H5 viruses, birds of the order *Charadriiformes* sporadically were found infected. Clade 2.3.4.4b of the Gs/Gd lineage was specifically successful in spreading to Europe and Africa since 2016. The lineage caused a major epidemic in white-winged terns (*Chlidonias leucopterus*) along the shores of Lake Victoria, Africa, in 2017 (15).

The 2020/2021 HPAI H5 epidemic in Europe surpassed all previously recorded European outbreaks in size, genotype constellations and reassortment frequency, with records of over 3,500 cases of lethally affected wild birds reported with laboratory-confirmed HPAIV H5 infection from 28 countries (16). Many ten-thousands of wild birds likely succumbed to infection without being virologically diagnosed. The 2.3.4.4b lineage proved to be highly promiscuous with respect to reassortment with cocirculating LPAIV. During the winter/spring season of 2020/2021 and continuing in 2022, several subtypes including H5N8, H5N1, H5N5, H5N4, and H5N3, and more than 30 genotypes have been identified in Europe (17, 18). While most sub- and many genotypes affected a range of both poultry and wild bird species, a unique HPAI H5N3 reassortant virus caused unusually high mortality rates only within a specific migratory wading bird niche, almost exclusively affecting red knots.

Here, we describe the brief but disastrous epidemic of a HPAIV clade 2.3.4.4b H5N3 reassortant in December 2020.

## Methods

Dead birds at the Schleswig-Holstein Wadden Sea National Park were collected and documented by rangers. While the birds were safely disposed by veterinary authorities, a batch of 513 individual red knot carcasses collected in December 2020 and kept frozen until August 2021 when they were dissected and assessed macroscopically under Biosafety Level 3 at the Friedrich-Loeffler-Institut, Isle of Riems in Germany.

Organ samples and combined oropharyngeal-cloacal swabs for virological analyses were collected during post mortem examination of ten randomly selected red knot carcasses representing 2% of the total of 513 dissected red knots. RNA-extraction from homogenized tissue material and swab sample medium was carried out as described by (19). Initially all RNA samples were tested for avian influenza virus genome applying generic, reverse transcription real-time PCR (RT-qPCR) targeting the influenza A virus nucleoprotein (NP), including an internal control assay (20, 21). In a subsequent step for each individual carcass, one RNA-sample with the lowest cycle threshold value (Ct value) in NP-qPCR was applied for further sub- and pathotype analyses using further previously described RT-qPCRs (RITA; (22)). The limit of detection of the subtyping PCRs is similar to that of the generic RT-qPCRs used and ranges between 25 to 250 per reaction (22).

To pinpoint the virus target tissue and cell tropism, immunohistochemistry (IHC) was performed for virus antigen detection using a primary antibody against the matrix1-protein of Influenza A virus (ATCC clone HB-64) as specified and the distribution of viral antigen was recorded in an ordinal scoring scale (score 0-4) as described by (23).

Viral full-genome sequences were generated from three red knots collected in Germany-Schleswig-Holstein and one red knot from Germany, Lower Saxony. Full-genome sequencing of AIV-positive samples was executed by a previously described nanopore-based real-time sequencing method with prior full genome amplification (24). For this, RNA extraction with the Qiagen Mini Viral Kit (Qiagen, Germany) and subsequent genome amplification with universal AIV-End-RT-PCR using Superscript III One-Step and Platinum Taq (ThermoFisher Scientific, USA) with one primer pair (Pan-IVA-1F: TCCCAGTCACGACGTCGTAGCGAAAGCAGG; Pan-IVA-1R: GGAAACAGCTATGACCATGAGTAGAAACAAGG), binding to the conserved ends of the AIV genome segments, was conducted. After purification of the PCR products with AMPure XP Magnetic Beads (Beckman-Coulter, USA), full-genome sequencing on a MinION platform (Oxford Nanopore Technologies, ONT, UK) using Rapid Barcoding Kit (SQK-RBK004, ONT) for transposon-based library preparation and multiplexing was performed. Sequencing was directed according to the manufacturer’s instructions with a R9.4.1 flow cell on Mk1C device with MinKNOW Software Core (v4.3.11). Live basecalling of the raw data with Guppy (v5.0.13, ONT) was followed by a demultiplexing, quality check and trimming step to remove low quality, primer and short (<50bp) sequences. After sequencing, full-genome consensus sequences were generated in a map-to-reference approach utilizing MiniMap2 (25). Reference genomes are a curated collection of all HA and NA subtypes alongside an assortment of internal gene sequences chosen to cover all potentially circulating viral strains. Polishing of the final genome sequences was done manually after consensus production according to the highest quality (60%) in Geneious Prime (v2021.0.1, Biomatters, New Zealand). For phylogenetic analysis sequences from EpiFlu^TM^ were retrieved where search was restricted to clade 2.3.4.4b H5N3 sequences or Eurasian non-GS/GD collected June 2019-May 2021. Respective accession numbers and data source acknowledgement can be found in Supplementary table S1.

Segment specific multiple alignments were generated using MAFFT (v7.450) (26) and subsequent maximum likelihood (ML) trees were calculated with RAxML (v8.2.11) (27) utilizing model GTR GAMMA with rapid bootstrapping and search for the best scoring ML tree supported with 1000 bootstrap replicates. Time-scaled trees of concatenated genomes of the same H5N3 genotype were calculated with the BEAST (v1.10.4) software package (28) using a GTR GAMMA substitution model, an uncorrelated relaxed clock with a lognormal distribution and coalescent constant population tree models. Chain lengths were set to 10 million iterations and convergence checked via Tracer (v1.7.1). Time-scaled summary maximum clade credibility trees (MCC) with 10% post burn-in posterior were created using TreeAnnotator (v1.10.4) and visualized with FigTree (V1.4.4). Robustness of the MCC trees was evaluated using 95% highest posterior density (HPD) confidence intervals at each node and posterior confidence values as branch support.

## Results

### Mass die-off of red knots

In Germany, more than 16,000 deceased or moribund waders and waterfowl had been identified in the Wadden Sea area of Schleswig-Holstein (predominantly in the district of Nordfriesland) between 25.October 2020 and end of March 2021 (fig. 1). Among all avian species, the highest number of HPAI cases was found in Barnacle geese (*Branta leucopsis*; 46%), red knots (21%) and Eurasian wigeons (*Mareca penelope*;10%) (fig. 1-2). While other species were found dead in high numbers throughout longer periods, 3,329 red knots were found dead mainly within only a short period between 14 to 23 December 2020 (fig. 2-3).

**Fig 1:**
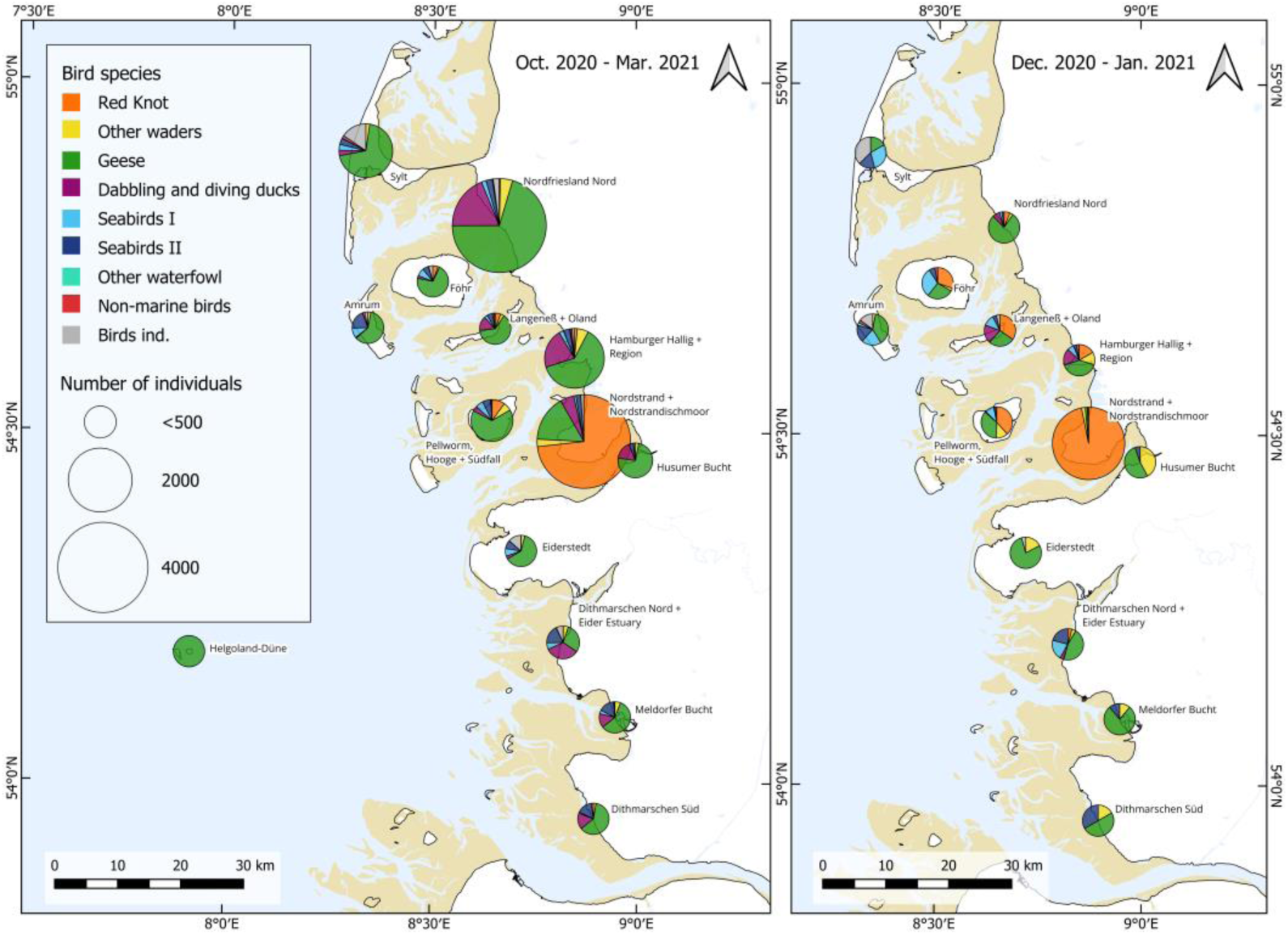
Location and species composition of collected dead birds in the period October 2020 to end of March 2021 (above) / during the mass die-off (15 December 2020 to 15 January 2021) in the northern German Wadden Sea. Note: red knots are marked in orange. Seabirds I: seaducks, mergansers, grebes, tubenoses, gannets, cormorants, auks. Seabirds II: gulls, skuas, terns.

**Fig 2:**
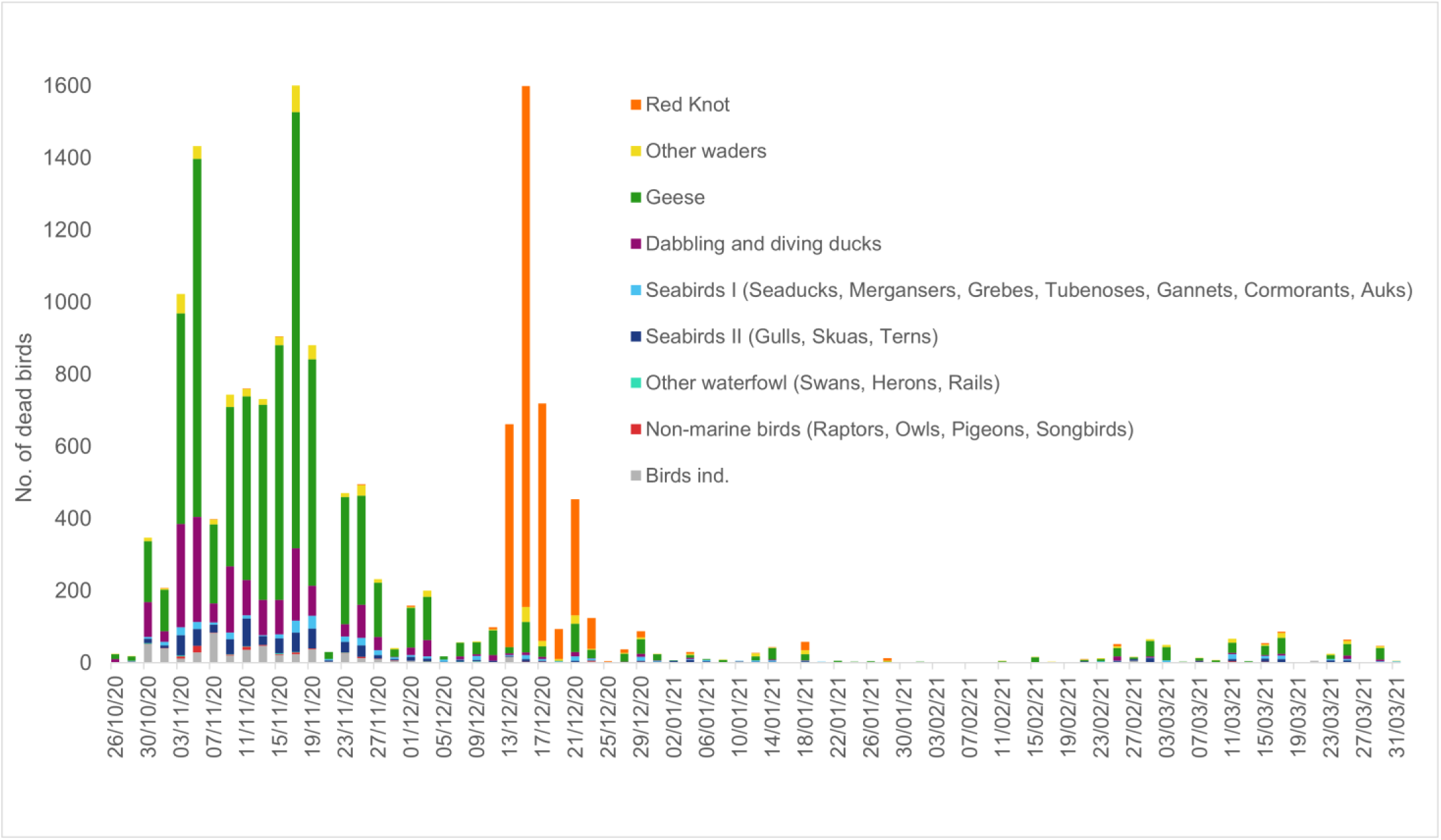
Number of dead birds allocated to different bird groups recorded in the Schleswig-Holstein Wadden Sea area within the period October 2020 to end of March 2021.

**Fig 3:**
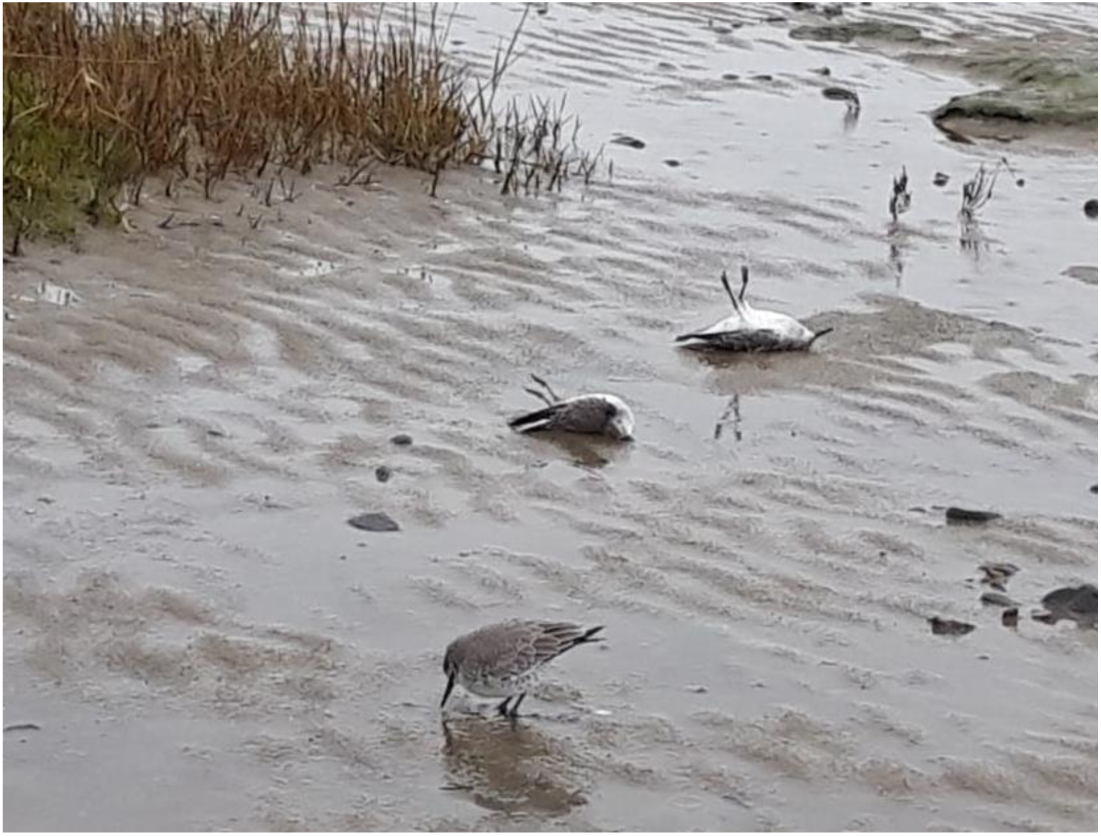
Red knots dead or dying at the high-tide roost (photo: Ruth Hartwig-Kruse)

The mass die-off in red knots occurred at a high tide roost on the peninsula of Nordstrand in the federal state of Schleswig-Holstein, Germany (54° 29' 15'' N; 8° 49' 04'' E; Fig. 1). Although both subspecies of the red knot cannot be distinguished securely morphologically, the virus outbreak described in this study affected exclusively individuals of the *islandica* subspecies as it took place in late December and therefore affected wintering birds. Most of the birds were already dead on the beach, while a few others were still able to fly and suddenly dropped dead from the sky to the ground. Other individuals were apathic and showed clear signs of neurological disorder with no escape reactions (fig 3). In addition to red knots only a common buzzard in the same region and time was detected with the same virus sub- and genotype, HPAIV H5N3. All other dead water birds were mainly HPAIV H5N8 positive at that time.

### Necropsy

A batch of 513 individual red knot carcasses was dissected and assessed macroscopically. Of these around 80% were adults and 20% were juveniles in their first calendar year. The sex distribution was uniform among adults and juveniles, respectively (overall 49 % of females and 51% of males were found). The macroscopic inspection of the 513 individuals showed a good to moderate body condition (according to van Franeker et al. 2007). A common finding was a nearly empty gut and a varying chest muscle thickness. In none of the individuals, the gut showed macroscopic lesions, whereas there were 10 (2%) individuals showing lesions of the liver. Furthermore, a total of 334 individuals (65%) showed internal bleedings in the lung and another total of 278 individuals (54%) had kidneys that were coloured whitish or were covered with white dots.

### Virologic and immunohistochemical results

As shown in table 1 influenza A virus genome was detected in every tested individual bird in at least eight out of nine tissue samples collected. Ct values ranged from 12.8 to 38.5. All individuals were confirmed to harbour HPAIV H5N3 RNA. The most prominent viral loads were detected in brain samples. Consistently, viral loads were also found in all ten oropharyngeal/cloacal swab samples, assuming virus shedding, although no virus isolation was conducted. In accordance with virus genome detection data, the brain was consistently affected (10 out of 10 birds). Further, the air sacs (8/10), the heart (7/10), and the kidneys (6/10) regularly exhibited virus antigen. Individual animals (1 or 2/10) showed virus protein labelling in the liver, lung, pancreas, large intestine, and skeletal muscle. No antigen was found in the proventriculus, gizzard and small intestine. The identified target cells comprised predominantly tissue-specific epithelial cells (parenchyma), however, some red knots presented with focal endotheliotropism or infection of smooth muscle cells. Details on target cells and antigen distribution are given in table 2, representative tissue slides are given in figure 4.

**Fig 4:**
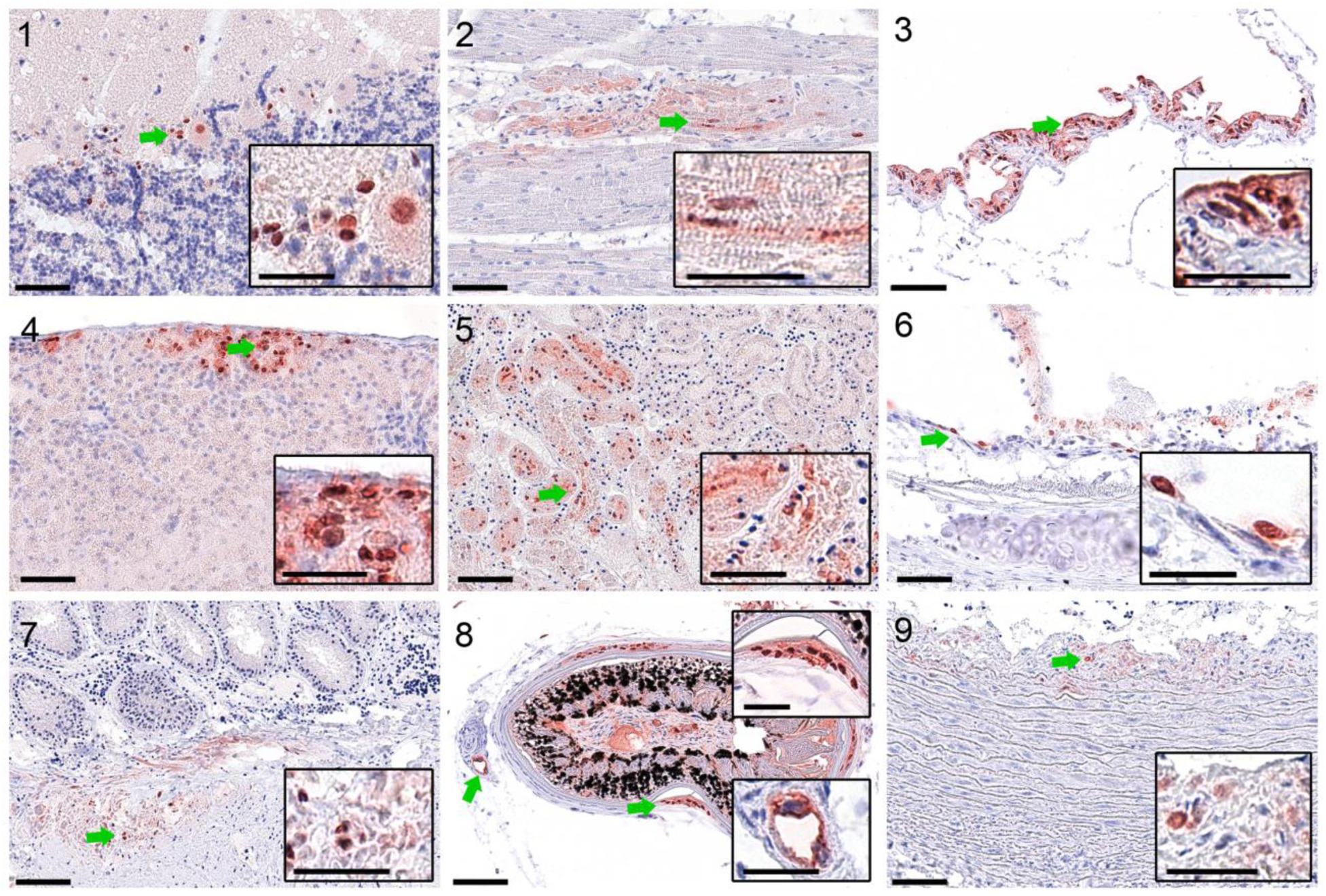
Influenza virus target cell tropism in red knots. Shown is the Influenza A matrix-1 protein (bright red staining) in neurons and glial cells, brain (1); cardiomyocytes, heart (2); epithelium, air sacs (3); hepatocytes, liver (4); tubular epithelium, kidney (5); bronchial epithelium, lung (6); smooth muscle cells, cecum (7); feather follicle epithelium and vascular endothelium, skin (8); smooth muscle cells, artery (9). Green arrows indicate the region of interest depicted in the inlay. Scale bars indicate 50 µm (original) or 25µm (inlays).

**Table 1a:** Distribution of viral genomic load based on NP2-RT-qPCR resp. H5-RT-qPCR in organ samples and/or swab samples of red knots found dead in Germany.

| Red knot carcasses |  |  |  |  |  |  |  |  |  |  |
| --- | --- | --- | --- | --- | --- | --- | --- | --- | --- | --- |
| Tissue sample | 1 | 2 | 3 | 4 | 5 | 6 | 7 | 8 | 9 | 10 |
| **Brain** | + | + | +++ | +++ | +++ | +++ | +++ | + | +++ | +++ |
| **Heart** | (+) | + | +++ | +++ | +++ | ++ | ++ | + | + | + |
| **Kidney** | (+) | + | +++ | +++ | ++ | ++ | ++ | + | +++ | (+) |
| **Liver** | n.a. | (+) | ++ | (+) | ++ | (+) | (+) | (+) | + | (+) |
| **Lung** | (+) | + | ++ | ++ | ++ | ++ | ++ | ++ | ++ | + |
| **Duodenum** | (+) | + | + | + | + | + | + | (+) | (+) | (+) |
| **Laying guts** | (+) | ++ | +++ | ++ | ++ | n.e. | ++ | n.e. | +++ | + |
| **Feather follicle** | (+) | (+) | + | - | (+) | n.e. | (+) | - | - | - |
| **Oph-cl swab** | (+) | (+) | + | + | + | + | (+) | + | + | (+) |

Influenza A virus generic RT-qPCR results of red knots no. 1-10 shown as Ct values: -, $\geq 40$ (negative); (+), $\geq 30-40$ ; +, $\geq 25-30$ ; ++, $\geq 20-25$ ; +++, $\leq 20$ . Red print indicates samples chosen for HP H5 subtype-confirmation. Liver sample of red knot no. 1 was not analysable. Feather sample of red knot no. 6 was not examined. Sex: #6, #8 male, all other birds female.

**Table 1b:** Sub- and pathotyping of influenza A virus RNA detected in brain samples of ten red knot carcasses collected in Germany.

| Red knot carcasses |  |  |  |  |  |  |  |  |  |  |
| --- | --- | --- | --- | --- | --- | --- | --- | --- | --- | --- |
| RT-qPCR | 1 | 2 | 3 | 4 | 5 | 6 | 7 | 8 | 9 | 10 |
| **H5** | ++ | ++ | +++ | +++ | +++ | +++ | ++ | ++ | +++ | +++ |
| **N3** | ++ | ++ | +++ | +++ | +++ | +++ | ++ | ++ | +++ | +++ |
| **HP-2.3.4.4b** | ++ | ++ | ++ | +++ | +++ | +++ | ++ | ++ | +++ | +++ |

RT-qPCR results for further sub- and pathotyping from chosen IAV-positive samples. Ct values are shown as described in Tab. 1a

**Table 2:** Virus antigen detection in tissues from ten red knot carcasses collected in Germany. Immunohistochemistry against Influenza matrix1-protein. - = negative, + = focal, ++ = multifocal, +++ = coalescent, ++++ = diffuse.

| Virus target tissues and cells in Red knots |  |  |  |  |  |  |  |  |  |  |  |
| --- | --- | --- | --- | --- | --- | --- | --- | --- | --- | --- | --- |
| Tissue | Target cells | 1 | 2 | 3 | 4 | 5 | 6 | 7 | 8 | 9 | 10 |
| Brain | neurons, glial cells, ependymal cell, choroid plexus epithelium | ++ | ++ | +++ | +++ | ++ | ++ | ++ | ++ | ++ | ++ |
| Heart | cardiomyocytes | - | + | ++ | +++ | ++ | ++ | +++ | + | - | - |
| Air sac | epithelium, luminal debris | - | + | +++ | + | + | + | ++ | + | + | - |
| Kidney | tubular epithelium | - | - | ++ | +++ | +++ | + | + | - | +++ | - |
| Liver | hepatocytes | - | - | + | - | + | - | - | - | - | - |
| Lung | bronchial epithelium | - | - | - | - | ++ | - | - | - | ++ | - |
| Pancreas | acinar epithelium | - | - | + | + | - | - | - | - | - | - |
| Proventriculus/<br>gizzard | - | - | - | - | - | - | - | - | - | - | - |
| Small intestine<br>(duodenum) | - | - | - | - | - | - | - | - | - | - | - |
| Large intestine,<br>cecum) | smooth muscle | - | - | + | - | - | + | - | - | - | - |
| Skeletal muscle | myocyte | - | - | - | + | - | - | - | - | - | - |
| Skin | feather follicle | - | - | + | - | - | - | - | + | - | - |
| Blood vessel,<br>any tissue | endothelium | - | - | + | - | - | - | + | - | - | - |
| Blood vessel,<br>any tissue | vessel, smooth muscle | - | - | - | + | - | - | - | - | - | - |

### Phylogenetic analysis

The goal of our phylogenetic analysis was to reconstruct the evolutionary relationship of HPAI H5 viruses, to identify common ancestors and the patterns of divergence over time. The search of clade 2.3.4.4b H5N3 sequences in the phylogenetic databases yielded a total of 13 genome sequences comprising HPAIV from “red knots” (n=5), “knot waders” (n=2), “Eurasian curlew (*Numenius arquata*)” (n=1), “common buzzard (*Buteo buteo*)” (n=2), “peregrine falcon (*Falco peregrinus*)” (n=2), and “common kestrel (*Falco tinnunculus*)” (n=1). Samples were collected in Denmark (n=1, “common kestrel”), France (n=2, “red knot”, “curlew”), Ireland (n=2, “knot wader”), Northern-Ireland (n=2, “peregrine falcon”), The Netherlands (n=2, “common buzzard”) and Germany (n=4, “red knots”). Exact sample details can be found in Table S1.

All respective H5N3 viruses belong to the H5 clade 2.3.4.4b while carrying a polybasic hemagglutinin cleavage site (HACS; PLREKRRKRGLF), thus fulfilling the legal criteria for high pathogenicity. Both the HA and matrix protein (MP) are highly similar to the previously described lineage of HPAI H5N8 viruses circulating in Central Europe from October 2020 onwards (29). All further six segments point towards reassortment with Eurasian avian lineage LPAIV genes as indicated by phylogenetic analyses of each the eight segments from a broad set of viruses covering geographic locations of Eurasia affected by HPAI (Supplementary Material 1-8; for the NA-gene, only NA3 genes were considered). The results are summarized in Figure 5. Closest relatives of the polymerase segments (polymerase basic 2 (PB2), polymerase basic 1 (PB1) and polymerase acidic (PA), were found in LPAI from wild birds, poultry, environmental and mixed samples across Europe from fall 2020 to early 2021 (PB2 and PB1 in A/turkey/Germany-BB/AI00868/2021, A/mallard/France/20P017917/2020, A/turkey/England/018179/2021; PA in A/environment/England/030642/2020). The NP segment was previously detected in a LPAIV H5N8 identified in Germany in September 2020 (A/guinea fowl/Germany-NW/AI01184/2020). The neuraminidase (NA), nucleoprotein (NP) and non-structural (NS) segments are most closely related to LPAIV genes detected in Europe over the past few years (A/mallard/France/20P017917/2020 (H5N3), A/turkey/England/018179/2021 (H5N3); NS in A/mallard/Denmark/12946-11/2020 (H7N5) A/Anas_platyrhynchos/Belgium/10413_0003/2020 (H5N2)). The H5N3 reassortant was one of several reassortments events resulting in different HPAIV subtypes (17).

**Fig 5:**
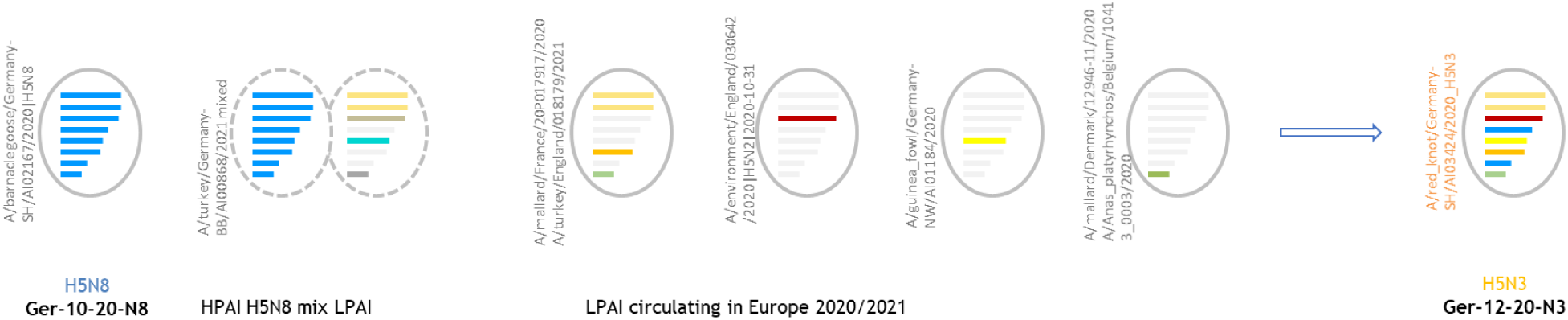
Visual representation of likely reassortment pathways of HPAIV clade 2.3.4.4b and Eurasian avian lineage LPAI viruses resulting in HPAI H5N3 viruses found in red knots in Europe. Dashed lines indicate mixed samples. Virus names of the closest relatives are given. The underlying trees can be found in Supplemental material 1-8. HPAI H5N3 of the red knots are composed of genes (in colour) of different viruses (from top to bottom: PB2, PB1, PA, HA, NP, NA, M, NS).

Segment-wise phylogenetic analyses of a broad set of sequences representative of different host species and geographical locations affected by HPAI H5 viruses indicate, in addition to high similarities over the entire genome range between 99.85 and 99.55% identity, a singular reassorting event generating the common ancestor of all HPAI H5N3 viruses from Germany analysed here. This timeframe fits to the result of the Bayesian phylogenetic inference analyses where the ancestral virus was calculated to have emerged in early November 2020.

## Discussion

The current study describes the first outbreak of a new reassortment of an HPAI virus subtype H5N3 of clade 2.3.4.4b causing a mass mortality in December 2020 in a group of *Charadrii* bird species in the German Wadden Sea that was never affected before in such high numbers. Mass die-offs of various bird species in the Wadden Sea have been reported before and were mainly attributed to food shortages (30, 31) and effects of severe weather (30, 32). Large botulism outbreaks occurred in the Elbe estuary in the 1980s and mid-1990s (33). In 2019, *Vibrio cholerae* was identified as a cause of high chick mortality here (34). In winter 2016/17, a first major avian influenza outbreak in wild birds (mainly wigeons) was detected in the Dutch Wadden Sea (35). Otherwise, monitoring of dead birds in the Wadden Sea in the past years have revealed a low degree (incidences <5%) of avian influenza infection (36).

The flyway population of red knots of the subspecies *islandica* is estimated 310,000 – 360,000 individuals (2) and shows a long-term declining trend which is currently recovering slowly since 2015 (5). The mortality described in this study affected around 3,500 red knots (which must be regarded as a minimum number given that likely not all dead individuals were found and accessible to counting) in the federal state of Schleswig-Holstein in Germany alone. Hence, the mass mortality event impacted around 1.0 - 1.1% of the whole flyway population. Exact numbers of red knots spending the winter in the Wadden Sea of the federal state of Schleswig-Holstein are unknown, but are estimated several ten thousand individuals (K. Günther, pers. comm.). Thus, the affected birds likely comprise a significant proportion of the population of wintering red knots in the Wadden Sea.

More than 500 red knots were subsequently dissected. No distinct pathological findings were present and birds seemed to be in good condition which may indicate a peracute course of the fatal disease which also was described earlier in an experimental infection with a HPAI H5 virus in red knots (37). A randomly selected number of samples taken for virological examinations yielded exclusively positive results indicating systemic HPAIV H5N3 infection. Accordingly, it can be assumed that all the red knots died of that infection. Furthermore, contemporary reports of mortalities of red knots being positive for HPAIV H5N3 in the Netherlands, France and the UK indicates a supraregional epidemic although significantly smaller numbers of red knots were impacted.

Red knots are known to use high tide roosts together with other *Charadriiformes* (6). At the same time and location other species, in particular barnacle geese, but in smaller numbers also dunlins and curlews were found dying from HPAIV infection but they tested positive for other H5 subtypes, mostly HPAIV H5N8. The HPAIV H5N3 reassortant was almost exclusively found in the red knots with the exception of buzzards, falcons and a kestrel that are assumed to have scavenged on the infected red knots. A single reassortment pathway involving gene segments from different HPAI H5 and LPAI viruses generated the common ancestor of all analysed HPAI H5N3 viruses from Germany. It was calculated that the ancestral virus emerged in early November 2020 and subsequently spread supra-regionally to other parts of northern and western Europe, but obviously on a smaller scale. After this event, this specific reassortant has not reappeared.

Red knots form dense social groups, both on their high tide roosts as well within their foraging sites (38). This might have facilitated intra-specific infection and restricted spread to other species. Other wader species such as oystercatchers (*Haematopus ostralegus*) that were largely spared from the epidemic are known to seek out (and defend) their own foraging sites on the intertidal mudflats separate from red knots (39–41).

Former studies have shown that red knots in the Wadden Sea are highly mobile and may switch between foraging sites and high tide roosts, respectively, in short periods of time (42, 43). In addition, virological investigations confirmed viral loads in oropharyngeal and cloacal samples suggesting virus shedding and transmission. Therefore, the question arises, why the virus did not spread to other places of the Wadden Sea and particularly to other high tide roosts in the vicinity to a greater extent. High virulence of the HPAIV H5N3 reassortant might provide an explanation: A peracute onset of disease with prominent neurological disorders as judged by the massive viral infection in brain tissues and reported clinical signs could have immobilized the birds rapidly whereby avoiding significant spread to other places in the Wadden Sea. Prominent loads of HPAIV H5N3-specific viral RNA were also found in heart muscle samples of dead red knots. Myocardial infection also would be in support of a hypothesis of rapid deaths as witnessed “birds falling dead from the sky”.

Former unrelated mass mortality events associated with starvation have shown that juvenile birds were impacted significantly more strongly than adults (32). Although the age composition of live red knots in the Wadden Sea during the year of the study is not known, the low proportion of juveniles among the birds found dead suggests that both age classes died in similar proportions. At least a high proportion of adults was concerned which is a crucial finding in terms of the impact on the overall population, as shorebirds are relatively long-lived and mortality of experienced adult birds is known to impact population dynamics significantly (44).

While the subtype HPAIV H5N3 seems to have completely vanished with the red knot fatalities, yet another subtype, HPAIV H5N1, emerged as the predominant virus on the Wadden Sea coast of Lower Saxony and Schleswig-Holstein since 2021, subsequently dominating all other infections in Germany, Europe and beyond. Since then, more than 4,600 cases of HPAIV H5N1 infection in wild birds have been reported from 34 EU countries, with Germany reporting the most cases in wild birds (EURL Avian Flu Data Portal izsvenezie.it). For the first time ever, the H5N1 subtype was still circulating during the summer months, with multiple introductions into breeding colonies of waterbird and seabird species and subsequent mass mortalities (45). During these outbreaks, no cases in red knots were reported from Germany.

Red knot subspecies *Calidris canutus rufa* and *Calidris canutus roselaari* have been investigated serologically in the U.S. (Delaware Bay and Alaska) with high antibody abundance against the LPAIV subtypes H3, H4, H10 and H11 (46, 1). Serological investigations of red knots have also been carried out in the East Atlantic Flyway with positive findings, while virological investigations yielded mostly negative results (12). It is unknown whether the affected red knots may have been pre-exposed to AIV before they were infected with HPAIV H5N3. Since colony breeding seabirds or wintering populations of red knots seem to be highly susceptible to lethal HPAIV H5 infection, they are more likely to be the victim of a spill-over event from a yet unknown source of infection than a potential future HPAIV reservoir species. However, more information is needed on the potential for and magnitude of survival of *Charadrii* species, including *Scolopacidae* and *Sternidae*.

## Conclusion and outlook

Since 2020, the avifauna of the Wadden Sea has been affected by HPAI H5 clade 2.3.4.4b on a larger scale. Although the incidence of HPAIV H5N3 infections with fatal outcomes in probably around 1% of the entire flyway population of *islandica* red knots was a singular and self-limiting event, similar reassortments in *Charadrii* species cannot be excluded in the future.

A combination of emergence of new reassortants, prolonged infection transmissions and introduction into breeding colonies of shorebirds will likely have the potential to severely impact the bird populations breeding, wintering and resting in the Wadden Sea World Heritage Site. In fact, such mass mortalities due to yet another gs/GD HPAIV reassortant of subtype H5N1 have been described in summer of 2022 (47, 45). In order to assess the resilience and level of threat of rare species, like the red knot, serological studies are needed to provide evidence on natural immunity and to estimate survival rates.

More frequent opportunities for possible spill-over to scavenging mammals due to amassed presence of avian carcasses harboring high viral loads, increase the risk for adaptation of HPAIV H5 clade 2.3.4.4.b viruses to mammals including humans. Hunted mammals like foxes, raccoon dogs, stray cats etc. shall be serologically investigated for influenza H5-specific antibodies to explore the extent of spill-over events in the region.

Observations, bird census, collection of deceased birds and mammals and their investigations for influenza viruses is crucial in understanding the evolution of influenza viruses and depends on strong collaboration of ornithologists, conservationists, state veterinarians, virologists and decision makers. Persons (rangers, bird ringers) in contact with potentially infected birds should be vaccinated against human influenza strains to prevent reassortments. Precautionary measures avoiding human exposure during carcass removal are indispensable.

## Acknowledgments

We would like to acknowledge and thank Aline Maksimov, Silvia Schuparis, Ralf Henkel and Diana Parlow for their excellent technical assistance. We thank the rangers and members of the Nationalpark Administration Schleswig-Holstein for collecting the dead birds. Two anonymous reviewers have helped significantly to improve the manuscript which is much appreciated.

## Author contributions

The author contributions are as listed:

Conceptualization: MB, TK, JK, AGl, PS, BH

Methodology: BH, CW, PS, SG

Investigation: ABa, EH, AK, AP, TCH, JK, AG, CG, BH, CW

Visualization: PS, ABa, JK, AP, AG, ABr, KH

Funding acquisition: MB

Supervision: PS, AGl, TCH, MB

Writing – original draft: JK, PS, AGl, TCH, AG, AB, AP, BH

Writing – review & editing: All authors

## Data and materials availability

All sequencing data has been deposited in the GISAID EpiFlu^TM^ sequence database and can be found under the accession numbers listed in Table S1.

## Supplementary Materials

Table S1; Figures Supplementary Material 1-8

