## Supplementary for "Red knots in Europe - a dead end host species or a new niche for highly pathogenic avian influenza?"

Supplementary material:

Maximum likelihood (ML) phylogeny for all 8 gene segments of the HPAI H5N3 virus detected in red knots in the Wadden Sea area in 2020.

Data source: sequence information with virus name, accession and metadata. We acknowledge the listed originating and submitting laboratories for sharing their data

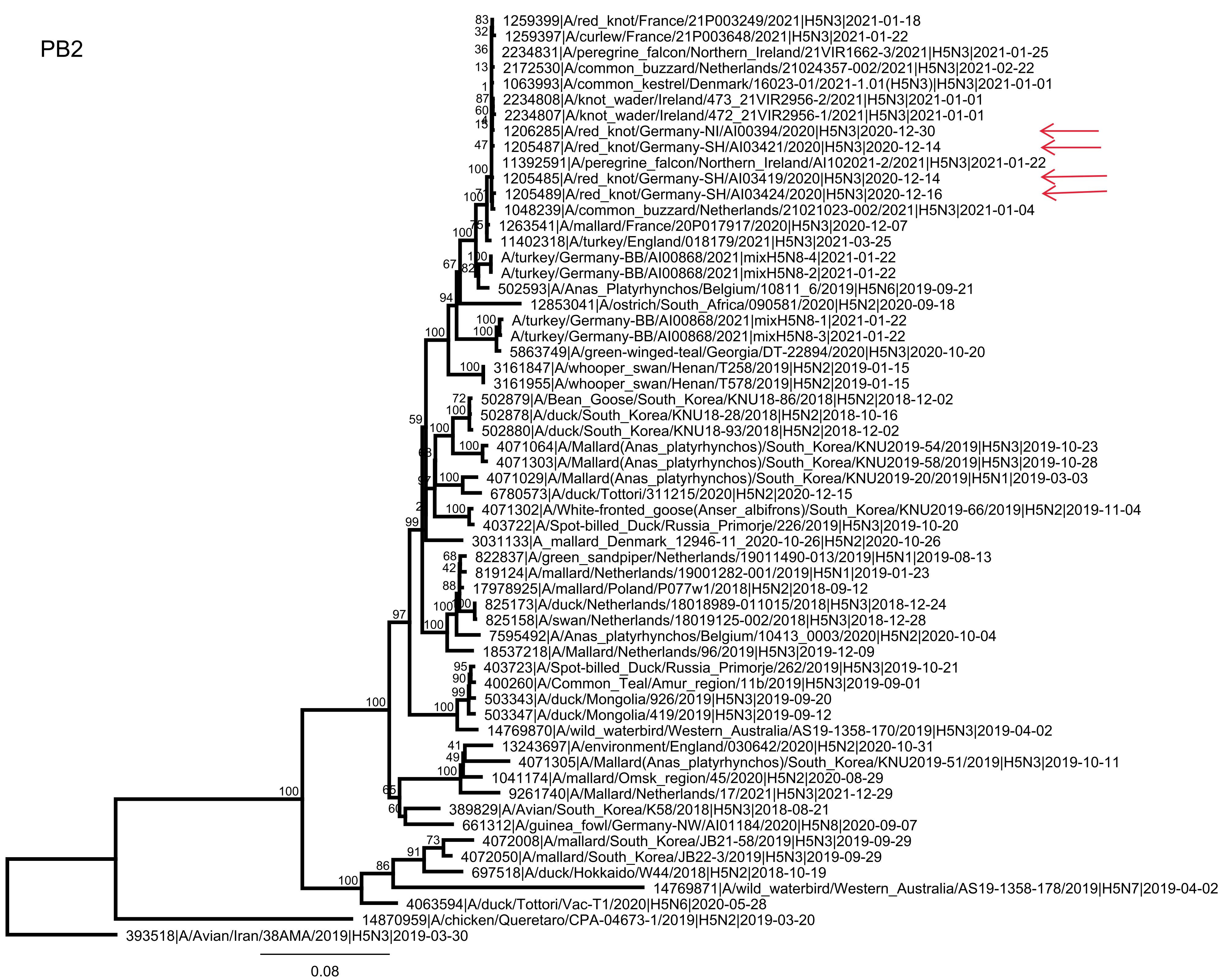
PB2
1259399|A/red_knot/France/21P003249/2021|H5N3|2021-01-18
1259397|A/curlew/France/21P003648/2021|H5N3|2021-01-22
2234831|A/peregrine_falcon/Northern_Ireland/21VIR1662-3/2021|H5N3|2021-01-25
2172530|A/common_buzzard/Netherlands/21024357-002/2021|H5N3|2021-02-22
1063993|A/common_kestrel/Denmark/16023-01/2021-1.01(H5N3)|H5N3|2021-01-01
2234808|A/knot_wader/Ireland/473_21VIR2956-2/2021|H5N3|2021-01-01
2234807|A/knot_wader/Ireland/472_21VIR2956-1/2021|H5N3|2021-01-01
1206285|A/red_knot/Germany-NI/AI00394/2020|H5N3|2020-12-30
1205487|A/red_knot/Germany-SH/AI03421/2020|H5N3|2020-12-14
11392591|A/peregrine_falcon/Northern_Ireland/AI102021-2/2021|H5N3|2021-01-22
1205485|A/red_knot/Germany-SH/AI03419/2020|H5N3|2020-12-14
1205489|A/red_knot/Germany-SH/AI03424/2020|H5N3|2020-12-16
1048239|A/common_buzzard/Netherlands/21021023-002/2021|H5N3|2021-01-04
1263541|A/mallard/France/20P017917/2020|H5N3|2020-12-07
11402318|A/turkey/England/018179/2021|H5N3|2021-03-25
A/turkey/Germany-BB/AI00868/2021|mixH5N8-4|2021-01-22
A/turkey/Germany-BB/AI00868/2021|mixH5N8-2|2021-01-22
502593|A/Anas_Platyrhynchos/Belgium/10811_6/2019|H5N6|2019-09-21
12853041|A/ostrich/South_Africa/090581/2020|H5N2|2020-09-18
A/turkey/Germany-BB/AI00868/2021|mixH5N8-1|2021-01-22
A/turkey/Germany-BB/AI00868/2021|mixH5N8-3|2021-01-22
5863749|A/green-winged-teal/Georgia/DT-22894/2020|H5N3|2020-10-20
3161847|A/whooper_swan/Henan/T258/2019|H5N2|2019-01-15
3161955|A/whooper_swan/Henan/T578/2019|H5N2|2019-01-15
502879|A/Bean_Goose/South_Korea/KNU18-86/2018|H5N2|2018-12-02
502878|A/duck/South_Korea/KNU18-28/2018|H5N2|2018-10-16
502880|A/duck/South_Korea/KNU18-93/2018|H5N2|2018-12-02
4071064|A/Mallard(Anas_platyrhynchos)/South_Korea/KNU2019-54/2019|H5N3|2019-10-23
4071303|A/Mallard(Anas_platyrhynchos)/South_Korea/KNU2019-58/2019|H5N3|2019-10-28
4071029|A/Mallard(Anas_platyrhynchos)/South_Korea/KNU2019-20/2019|H5N1|2019-03-03
6780573|A/duck/Tottori/311215/2020|H5N2|2020-12-15
4071302|A/White-fronted_goose(Anser_albifrons)/South_Korea/KNU2019-66/2019|H5N2|2019-11-04
403722|A/Spot-billed_Duck/Russia_Primorje/226/2019|H5N3|2019-10-20
3031133|A_mallard_Denmark_12946-11_2020-10-26|H5N2|2020-10-26
822837|A/green_sandpiper/Netherlands/19011490-013/2019|H5N1|2019-08-13
819124|A/mallard/Netherlands/19001282-001/2019|H5N1|2019-01-23
17978925|A/mallard/Poland/P077w1/2018|H5N2|2018-09-12
825173|A/duck/Netherlands/18018989-011015/2018|H5N3|2018-12-24
825158|A/swan/Netherlands/18019125-002/2018|H5N3|2018-12-28
7595492|A/Anas_platyrhynchos/Belgium/10413_0003/2020|H5N2|2020-10-04
18537218|A/Mallard/Netherlands/96/2019|H5N3|2019-12-09
403723|A/Spot-billed_Duck/Russia_Primorje/262/2019|H5N3|2019-10-21
400260|A/Common_Teal/Amur_region/11b/2019|H5N3|2019-09-01
503343|A/duck/Mongolia/926/2019|H5N3|2019-09-20
503347|A/duck/Mongolia/419/2019|H5N3|2019-09-12
14769870|A/wild_waterbird/Western_Australia/AS19-1358-170/2019|H5N3|2019-04-02
13243697|A/environment/England/030642/2020|H5N2|2020-10-31
4071305|A/Mallard(Anas_platyrhynchos)/South_Korea/KNU2019-51/2019|H5N3|2019-10-11
1041174|A/mallard/Omsk_region/45/2020|H5N2|2020-08-29
9261740|A/Mallard/Netherlands/17/2021|H5N3|2021-12-29
389829|A/Avian/South_Korea/K58/2018|H5N3|2018-08-21
661312|A/guinea_fowl/Germany-NW/AI01184/2020|H5N8|2020-09-07
4072008|A/mallard/South_Korea/JB21-58/2019|H5N3|2019-09-29
4072050|A/mallard/South_Korea/JB22-3/2019|H5N3|2019-09-29
697518|A/duck/Hokkaido/W44/2018|H5N2|2018-10-19
14769871|A/wild_waterbird/Western_Australia/AS19-1358-178/2019|H5N7|2019-04-02
4063594|A/duck/Tottori/Vac-T1/2020|H5N6|2020-05-28
14870959|A/chicken/Queretaro/CPA-04673-1/2019|H5N2|2019-03-20
393518|A/Avian/Iran/38AMA/2019|H5N3|2019-03-30
0.08

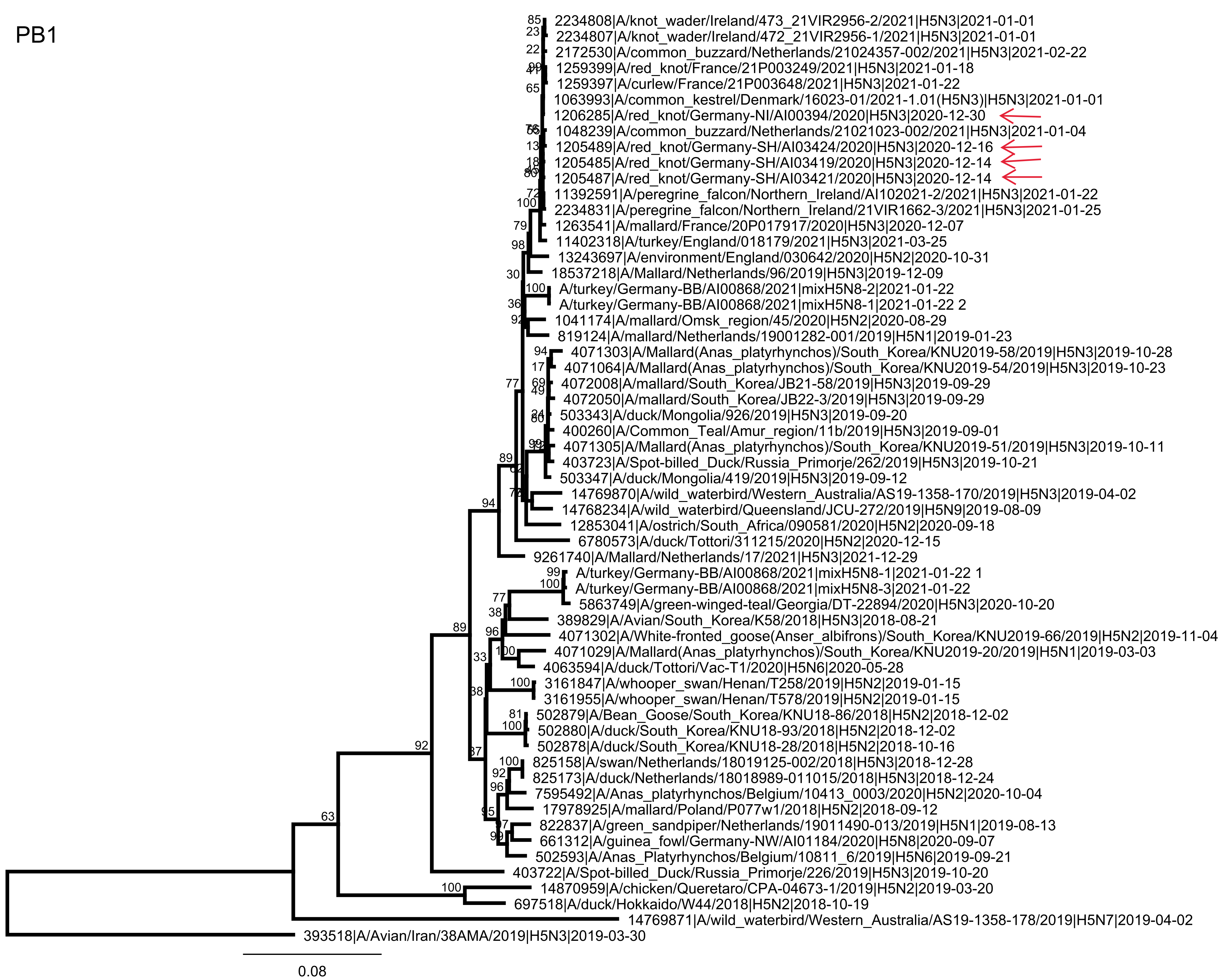
PB1
2234808|A/knot_wader/Ireland/473_21VIR2956-2/2021|H5N3|2021-01-01
2234807|A/knot_wader/Ireland/472_21VIR2956-1/2021|H5N3|2021-01-01
2172530|A/common_buzzard/Netherlands/21024357-002/2021|H5N3|2021-02-22
1259399|A/red_knot/France/21P003249/2021|H5N3|2021-01-18
1259397|A/curlew/France/21P003648/2021|H5N3|2021-01-22
1063993|A/common_kestrel/Denmark/16023-01/2021-1.01(H5N3)|H5N3|2021-01-01
1206285|A/red_knot/Germany-NI/AI00394/2020|H5N3|2020-12-30
1048239|A/common_buzzard/Netherlands/21021023-002/2021|H5N3|2021-01-04
1205489|A/red_knot/Germany-SH/AI03424/2020|H5N3|2020-12-16
1205485|A/red_knot/Germany-SH/AI03419/2020|H5N3|2020-12-14
1205487|A/red_knot/Germany-SH/AI03421/2020|H5N3|2020-12-14
11392591|A/peregrine_falcon/Northern_Ireland/AI102021-2/2021|H5N3|2021-01-22
2234831|A/peregrine_falcon/Northern_Ireland/21VIR1662-3/2021|H5N3|2021-01-25
1263541|A/mallard/France/20P017917/2020|H5N3|2020-12-07
11402318|A/turkey/England/018179/2021|H5N3|2021-03-25
13243697|A/environment/England/030642/2020|H5N2|2020-10-31
18537218|A/Mallard/Netherlands/96/2019|H5N3|2019-12-09
A/turkey/Germany-BB/AI00868/2021|mixH5N8-2|2021-01-22
A/turkey/Germany-BB/AI00868/2021|mixH5N8-1|2021-01-22 2
1041174|A/mallard/Omsk_region/45/2020|H5N2|2020-08-29
819124|A/mallard/Netherlands/19001282-001/2019|H5N1|2019-01-23
4071303|A/Mallard(Anas_platyrhynchos)/South_Korea/KNU2019-58/2019|H5N3|2019-10-28
4071064|A/Mallard(Anas_platyrhynchos)/South_Korea/KNU2019-54/2019|H5N3|2019-10-23
4072008|A/mallard/South_Korea/JB21-58/2019|H5N3|2019-09-29
4072050|A/mallard/South_Korea/JB22-3/2019|H5N3|2019-09-29
503343|A/duck/Mongolia/926/2019|H5N3|2019-09-20
400260|A/Common_Teal/Amur_region/11b/2019|H5N3|2019-09-01
4071305|A/Mallard(Anas_platyrhynchos)/South_Korea/KNU2019-51/2019|H5N3|2019-10-11
403723|A/Spot-billed_Duck/Russia_Primorje/262/2019|H5N3|2019-10-21
503347|A/duck/Mongolia/419/2019|H5N3|2019-09-12
14769870|A/wild_waterbird/Western_Australia/AS19-1358-170/2019|H5N3|2019-04-02
14768234|A/wild_waterbird/Queensland/JCU-272/2019|H5N9|2019-08-09
12853041|A/ostrich/South_Africa/090581/2020|H5N2|2020-09-18
6780573|A/duck/Tottori/311215/2020|H5N2|2020-12-15
9261740|A/Mallard/Netherlands/17/2021|H5N3|2021-12-29
A/turkey/Germany-BB/AI00868/2021|mixH5N8-1|2021-01-22 1
A/turkey/Germany-BB/AI00868/2021|mixH5N8-3|2021-01-22
5863749|A/green-winged-teal/Georgia/DT-22894/2020|H5N3|2020-10-20
389829|A/Avian/South_Korea/K58/2018|H5N3|2018-08-21
4071302|A/White-fronted_goose(Anser_albifrons)/South_Korea/KNU2019-66/2019|H5N2|2019-11-04
4071029|A/Mallard(Anas_platyrhynchos)/South_Korea/KNU2019-20/2019|H5N1|2019-03-03
4063594|A/duck/Tottori/Vac-T1/2020|H5N6|2020-05-28
3161847|A/whooper_swan/Henan/T258/2019|H5N2|2019-01-15
3161955|A/whooper_swan/Henan/T578/2019|H5N2|2019-01-15
502879|A/Bean_Goose/South_Korea/KNU18-86/2018|H5N2|2018-12-02
502880|A/duck/South_Korea/KNU18-93/2018|H5N2|2018-12-02
502878|A/duck/South_Korea/KNU18-28/2018|H5N2|2018-10-16
825158|A/swan/Netherlands/18019125-002/2018|H5N3|2018-12-28
825173|A/duck/Netherlands/18018989-011015/2018|H5N3|2018-12-24
7595492|A/Anas_platyrhynchos/Belgium/10413_0003/2020|H5N2|2020-10-04
17978925|A/mallard/Poland/P077w1/2018|H5N2|2018-09-12
822837|A/green_sandpiper/Netherlands/19011490-013/2019|H5N1|2019-08-13
661312|A/guinea_fowl/Germany-NW/AI01184/2020|H5N8|2020-09-07
502593|A/Anas_Platyrhynchos/Belgium/10811_6/2019|H5N6|2019-09-21
403722|A/Spot-billed_Duck/Russia_Primorje/226/2019|H5N3|2019-10-20
14870959|A/chicken/Queretaro/CPA-04673-1/2019|H5N2|2019-03-20
697518|A/duck/Hokkaido/W44/2018|H5N2|2018-10-19
14769871|A/wild_waterbird/Western_Australia/AS19-1358-178/2019|H5N7|2019-04-02
393518|A/Avian/Iran/38AMA/2019|H5N3|2019-03-30
0.08

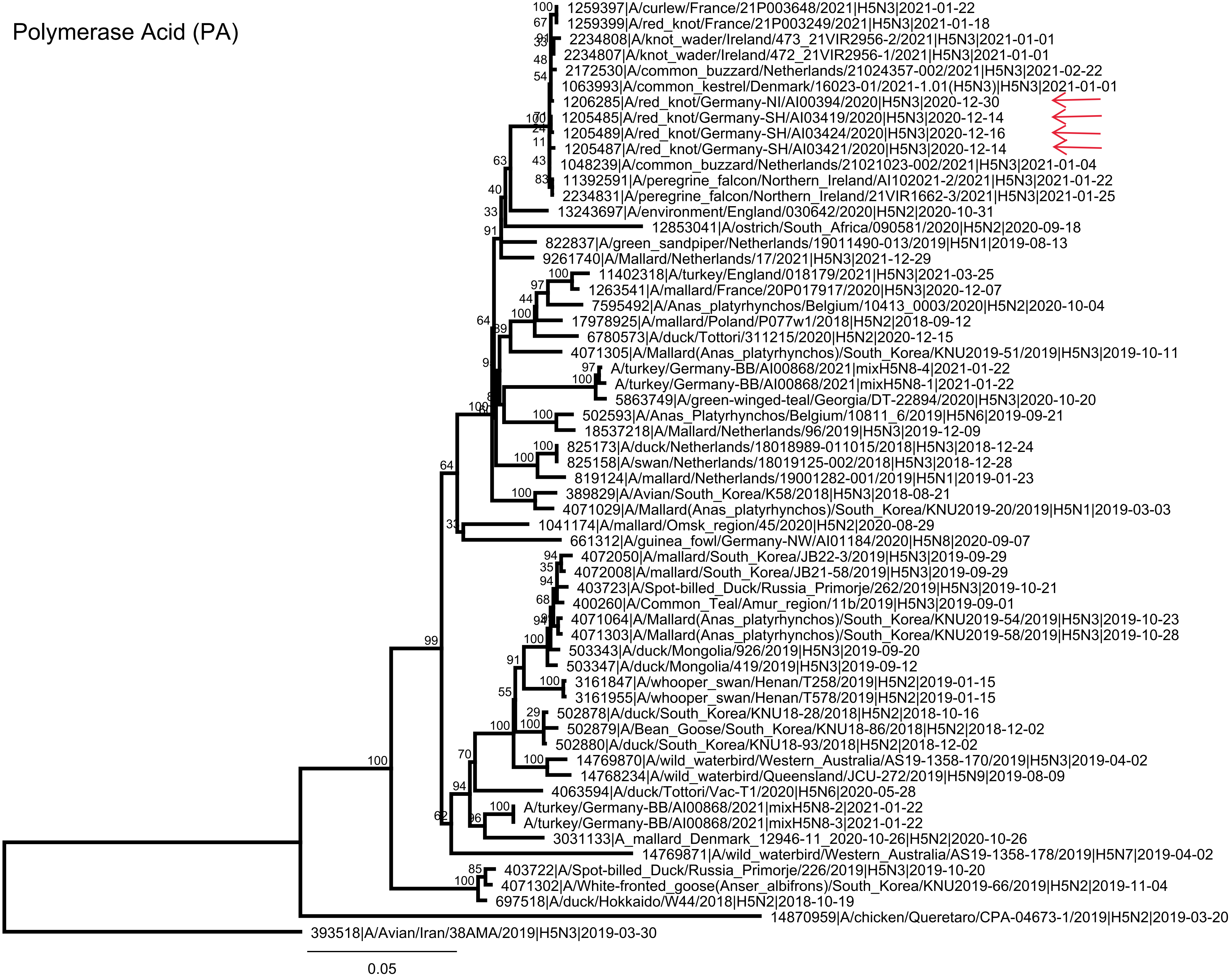
Polymerase Acid (PA)
1259397|A/curlew/France/21P003648/2021|H5N3|2021-01-22
1259399|A/red_knot/France/21P003249/2021|H5N3|2021-01-18
2234808|A/knot_wader/Ireland/473_21VIR2956-2/2021|H5N3|2021-01-01
2234807|A/knot_wader/Ireland/472_21VIR2956-1/2021|H5N3|2021-01-01
2172530|A/common_buzzard/Netherlands/21024357-002/2021|H5N3|2021-02-22
1063993|A/common_kestrel/Denmark/16023-01/2021-1.01(H5N3)|H5N3|2021-01-01
1206285|A/red_knot/Germany-NI/AI00394/2020|H5N3|2020-12-30
1205485|A/red_knot/Germany-SH/AI03419/2020|H5N3|2020-12-14
1205489|A/red_knot/Germany-SH/AI03424/2020|H5N3|2020-12-16
1205487|A/red_knot/Germany-SH/AI03421/2020|H5N3|2020-12-14
1048239|A/common_buzzard/Netherlands/21021023-002/2021|H5N3|2021-01-04
11392591|A/peregrine_falcon/Northern_Ireland/AI102021-2/2021|H5N3|2021-01-22
2234831|A/peregrine_falcon/Northern_Ireland/21VIR1662-3/2021|H5N3|2021-01-25
13243697|A/environment/England/030642/2020|H5N2|2020-10-31
12853041|A/ostrich/South_Africa/090581/2020|H5N2|2020-09-18
822837|A/green_sandpiper/Netherlands/19011490-013/2019|H5N1|2019-08-13
9261740|A/Mallard/Netherlands/17/2021|H5N3|2021-12-29
11402318|A/turkey/England/018179/2021|H5N3|2021-03-25
1263541|A/mallard/France/20P017917/2020|H5N3|2020-12-07
7595492|A/Anas_platyrhynchos/Belgium/10413_0003/2020|H5N2|2020-10-04
17978925|A/mallard/Poland/P077w1/2018|H5N2|2018-09-12
6780573|A/duck/Tottori/311215/2020|H5N2|2020-12-15
4071305|A/Mallard(Anas_platyrhynchos)/South_Korea/KNU2019-51/2019|H5N3|2019-10-11
A/turkey/Germany-BB/AI00868/2021|mixH5N8-4|2021-01-22
A/turkey/Germany-BB/AI00868/2021|mixH5N8-1|2021-01-22
5863749|A/green-winged-teal/Georgia/DT-22894/2020|H5N3|2020-10-20
502593|A/Anas_Platyrhynchos/Belgium/10811_6/2019|H5N6|2019-09-21
18537218|A/Mallard/Netherlands/96/2019|H5N3|2019-12-09
825173|A/duck/Netherlands/18018989-011015/2018|H5N3|2018-12-24
825158|A/swan/Netherlands/18019125-002/2018|H5N3|2018-12-28
819124|A/mallard/Netherlands/19001282-001/2019|H5N1|2019-01-23
389829|A/Avian/South_Korea/K58/2018|H5N3|2018-08-21
4071029|A/Mallard(Anas_platyrhynchos)/South_Korea/KNU2019-20/2019|H5N1|2019-03-03
1041174|A/mallard/Omsk_region/45/2020|H5N2|2020-08-29
661312|A/guinea_fowl/Germany-NW/AI01184/2020|H5N8|2020-09-07
4072050|A/mallard/South_Korea/JB22-3/2019|H5N3|2019-09-29
4072008|A/mallard/South_Korea/JB21-58/2019|H5N3|2019-09-29
403723|A/Spot-billed_Duck/Russia_Primorje/262/2019|H5N3|2019-10-21
400260|A/Common_Teal/Amur_region/11b/2019|H5N3|2019-09-01
4071064|A/Mallard(Anas_platyrhynchos)/South_Korea/KNU2019-54/2019|H5N3|2019-10-23
4071303|A/Mallard(Anas_platyrhynchos)/South_Korea/KNU2019-58/2019|H5N3|2019-10-28
503343|A/duck/Mongolia/926/2019|H5N3|2019-09-20
503347|A/duck/Mongolia/419/2019|H5N3|2019-09-12
3161847|A/whooper_swan/Henan/T258/2019|H5N2|2019-01-15
3161955|A/whooper_swan/Henan/T578/2019|H5N2|2019-01-15
502878|A/duck/South_Korea/KNU18-28/2018|H5N2|2018-10-16
502879|A/Bean_Goose/South_Korea/KNU18-86/2018|H5N2|2018-12-02
502880|A/duck/South_Korea/KNU18-93/2018|H5N2|2018-12-02
14769870|A/wild_waterbird/Western_Australia/AS19-1358-170/2019|H5N3|2019-04-02
14768234|A/wild_waterbird/Queensland/JCU-272/2019|H5N9|2019-08-09
4063594|A/duck/Tottori/Vac-T1/2020|H5N6|2020-05-28
A/turkey/Germany-BB/AI00868/2021|mixH5N8-2|2021-01-22
A/turkey/Germany-BB/AI00868/2021|mixH5N8-3|2021-01-22
3031133|A_mallard_Denmark_12946-11_2020-10-26|H5N2|2020-10-26
14769871|A/wild_waterbird/Western_Australia/AS19-1358-178/2019|H5N7|2019-04-02
403722|A/Spot-billed_Duck/Russia_Primorje/226/2019|H5N3|2019-10-20
4071302|A/White-fronted_goose(Anser_albifrons)/South_Korea/KNU2019-66/2019|H5N2|2019-11-04
697518|A/duck/Hokkaido/W44/2018|H5N2|2018-10-19
14870959|A/chicken/Queretaro/CPA-04673-1/2019|H5N2|2019-03-20
393518|A/Avian/Iran/38AMA/2019|H5N3|2019-03-30
0.05

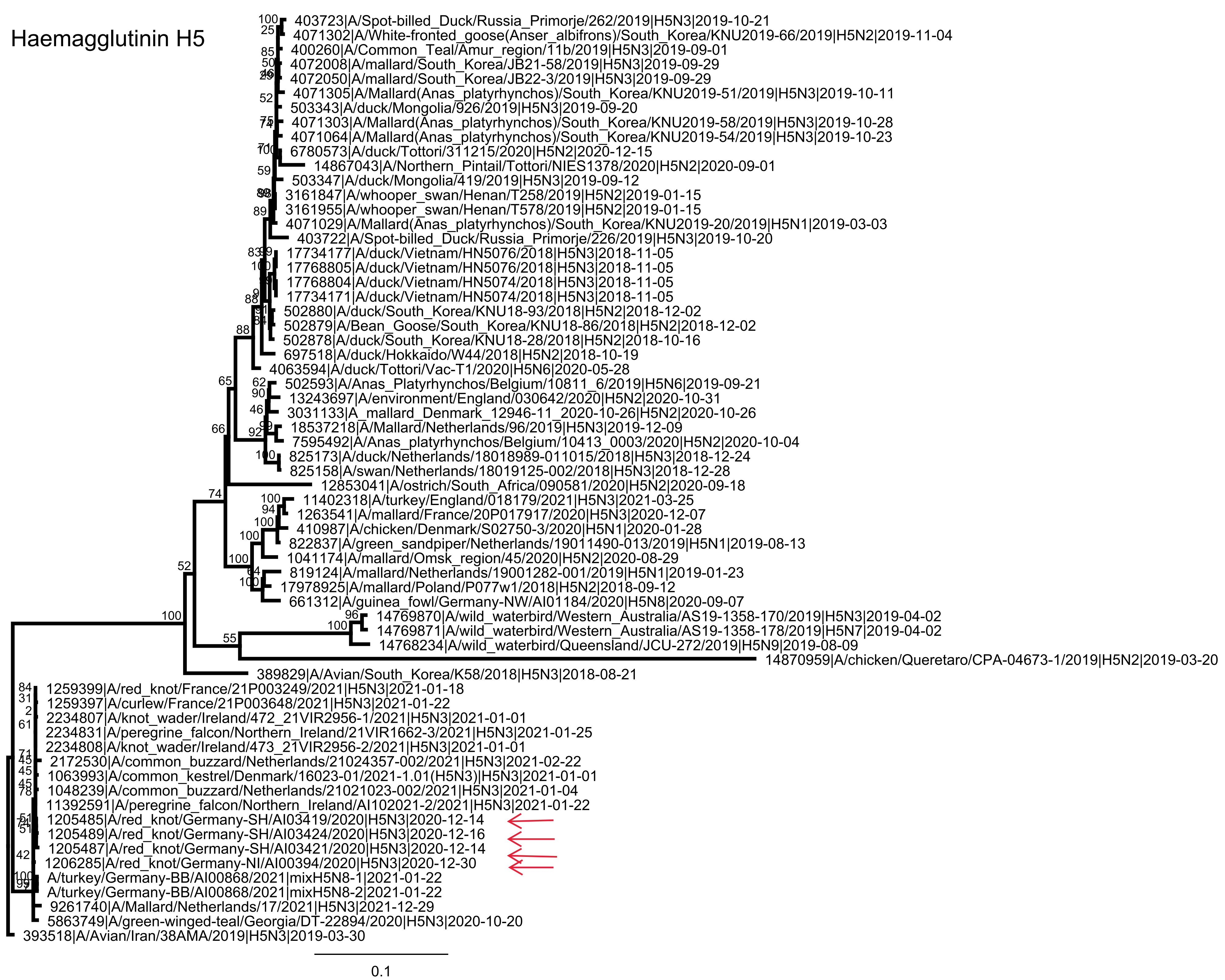
Haemagglutinin H5
403723|A/Spot-billed_Duck/Russia_Primorje/262/2019|H5N3|2019-10-21
4071302|A/White-fronted_goose(Anser_albifrons)/South_Korea/KNU2019-66/2019|H5N2|2019-11-04
400260|A/Common_Teal/Amur_region/11b/2019|H5N3|2019-09-01
4072008|A/mallard/South_Korea/JB21-58/2019|H5N3|2019-09-29
4072050|A/mallard/South_Korea/JB22-3/2019|H5N3|2019-09-29
4071305|A/Mallard(Anas_platyrhynchos)/South_Korea/KNU2019-51/2019|H5N3|2019-10-11
503343|A/duck/Mongolia/926/2019|H5N3|2019-09-20
4071303|A/Mallard(Anas_platyrhynchos)/South_Korea/KNU2019-58/2019|H5N3|2019-10-28
4071064|A/Mallard(Anas_platyrhynchos)/South_Korea/KNU2019-54/2019|H5N3|2019-10-23
6780573|A/duck/Tottori/311215/2020|H5N2|2020-12-15
14867043|A/Northern_Pintail/Tottori/NIES1378/2020|H5N2|2020-09-01
503347|A/duck/Mongolia/419/2019|H5N3|2019-09-12
3161847|A/whooper_swan/Henan/T258/2019|H5N2|2019-01-15
3161955|A/whooper_swan/Henan/T578/2019|H5N2|2019-01-15
4071029|A/Mallard(Anas_platyrhynchos)/South_Korea/KNU2019-20/2019|H5N1|2019-03-03
403722|A/Spot-billed_Duck/Russia_Primorje/226/2019|H5N3|2019-10-20
17734177|A/duck/Vietnam/HN5076/2018|H5N3|2018-11-05
17768805|A/duck/Vietnam/HN5076/2018|H5N3|2018-11-05
17768804|A/duck/Vietnam/HN5074/2018|H5N3|2018-11-05
17734171|A/duck/Vietnam/HN5074/2018|H5N3|2018-11-05
502880|A/duck/South_Korea/KNU18-93/2018|H5N2|2018-12-02
502879|A/Bean_Goose/South_Korea/KNU18-86/2018|H5N2|2018-12-02
502878|A/duck/South_Korea/KNU18-28/2018|H5N2|2018-10-16
697518|A/duck/Hokkaido/W44/2018|H5N2|2018-10-19
4063594|A/duck/Tottori/Vac-T1/2020|H5N6|2020-05-28
502593|A/Anas_Platyrhynchos/Belgium/10811_6/2019|H5N6|2019-09-21
13243697|A/environment/England/030642/2020|H5N2|2020-10-31
3031133|A_mallard_Denmark_12946-11_2020-10-26|H5N2|2020-10-26
18537218|A/Mallard/Netherlands/96/2019|H5N3|2019-12-09
7595492|A/Anas_platyrhynchos/Belgium/10413_0003/2020|H5N2|2020-10-04
825173|A/duck/Netherlands/18018989-011015/2018|H5N3|2018-12-24
825158|A/swan/Netherlands/18019125-002/2018|H5N3|2018-12-28
12853041|A/ostrich/South_Africa/090581/2020|H5N2|2020-09-18
11402318|A/turkey/England/018179/2021|H5N3|2021-03-25
1263541|A/mallard/France/20P017917/2020|H5N3|2020-12-07
410987|A/chicken/Denmark/S02750-3/2020|H5N1|2020-01-28
822837|A/green_sandpiper/Netherlands/19011490-013/2019|H5N1|2019-08-13
1041174|A/mallard/Omsk_region/45/2020|H5N2|2020-08-29
819124|A/mallard/Netherlands/19001282-001/2019|H5N1|2019-01-23
17978925|A/mallard/Poland/P077w1/2018|H5N2|2018-09-12
661312|A/guinea_fowl/Germany-NW/AI01184/2020|H5N8|2020-09-07
14769870|A/wild_waterbird/Western_Australia/AS19-1358-170/2019|H5N3|2019-04-02
14769871|A/wild_waterbird/Western_Australia/AS19-1358-178/2019|H5N7|2019-04-02
14768234|A/wild_waterbird/Queensland/JCU-272/2019|H5N9|2019-08-09
14870959|A/chicken/Queretaro/CPA-04673-1/2019|H5N2|2019-03-20
389829|A/Avian/South_Korea/K58/2018|H5N3|2018-08-21
1259399|A/red_knot/France/21P003249/2021|H5N3|2021-01-18
1259397|A/curlew/France/21P003648/2021|H5N3|2021-01-22
2234807|A/knot_wader/Ireland/472_21VIR2956-1/2021|H5N3|2021-01-01
2234831|A/peregrine_falcon/Northern_Ireland/21VIR1662-3/2021|H5N3|2021-01-25
2234808|A/knot_wader/Ireland/473_21VIR2956-2/2021|H5N3|2021-01-01
2172530|A/common_buzzard/Netherlands/21024357-002/2021|H5N3|2021-02-22
1063993|A/common_kestrel/Denmark/16023-01/2021-1.01(H5N3)|H5N3|2021-01-01
1048239|A/common_buzzard/Netherlands/21021023-002/2021|H5N3|2021-01-04
11392591|A/peregrine_falcon/Northern_Ireland/AI102021-2/2021|H5N3|2021-01-22
1205485|A/red_knot/Germany-SH/AI03419/2020|H5N3|2020-12-14
1205489|A/red_knot/Germany-SH/AI03424/2020|H5N3|2020-12-16
1205487|A/red_knot/Germany-SH/AI03421/2020|H5N3|2020-12-14
1206285|A/red_knot/Germany-NI/AI00394/2020|H5N3|2020-12-30
A/turkey/Germany-BB/AI00868/2021|mixH5N8-1|2021-01-22
A/turkey/Germany-BB/AI00868/2021|mixH5N8-2|2021-01-22
9261740|A/Mallard/Netherlands/17/2021|H5N3|2021-12-29
5863749|A/green-winged-teal/Georgia/DT-22894/2020|H5N3|2020-10-20
393518|A/Avian/Iran/38AMA/2019|H5N3|2019-03-30
0.1

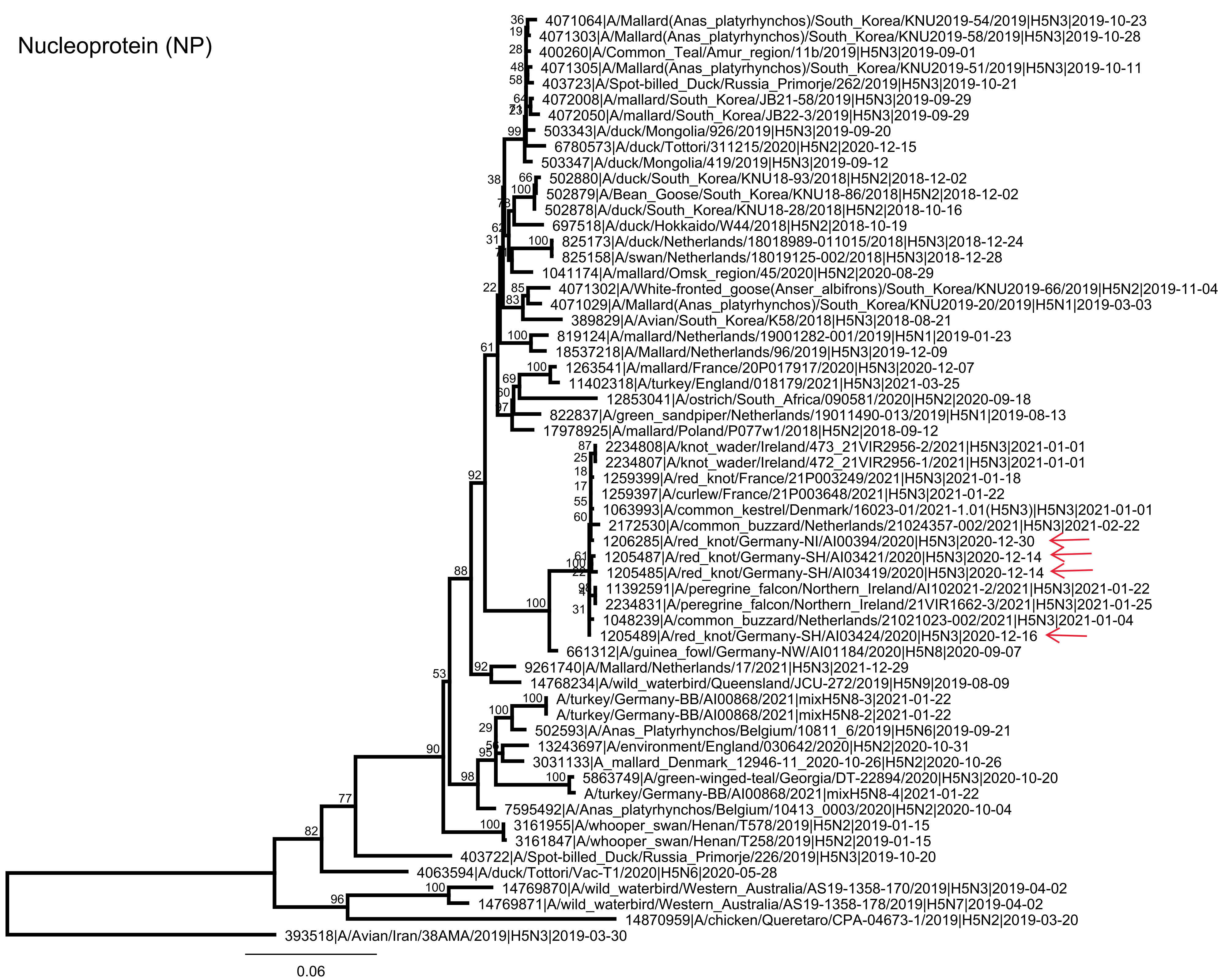

Nucleoprotein (NP)
4071064|A/Mallard(Anas_platyrhynchos)/South_Korea/KNU2019-54/2019|H5N3|2019-10-23
4071303|A/Mallard(Anas_platyrhynchos)/South_Korea/KNU2019-58/2019|H5N3|2019-10-28
400260|A/Common_Teal/Amur_region/11b/2019|H5N3|2019-09-01
4071305|A/Mallard(Anas_platyrhynchos)/South_Korea/KNU2019-51/2019|H5N3|2019-10-11
403723|A/Spot-billed_Duck/Russia_Primorje/262/2019|H5N3|2019-10-21
4072008|A/mallard/South_Korea/JB21-58/2019|H5N3|2019-09-29
4072050|A/mallard/South_Korea/JB22-3/2019|H5N3|2019-09-29
503343|A/duck/Mongolia/926/2019|H5N3|2019-09-20
6780573|A/duck/Tottori/311215/2020|H5N2|2020-12-15
503347|A/duck/Mongolia/419/2019|H5N3|2019-09-12
502880|A/duck/South_Korea/KNU18-93/2018|H5N2|2018-12-02
502879|A/Bean_Goose/South_Korea/KNU18-86/2018|H5N2|2018-12-02
502878|A/duck/South_Korea/KNU18-28/2018|H5N2|2018-10-16
697518|A/duck/Hokkaido/W44/2018|H5N2|2018-10-19
825173|A/duck/Netherlands/18018989-011015/2018|H5N3|2018-12-24
825158|A/swan/Netherlands/18019125-002/2018|H5N3|2018-12-28
1041174|A/mallard/Omsk_region/45/2020|H5N2|2020-08-29
4071302|A/White-fronted_goose(Anser_albifrons)/South_Korea/KNU2019-66/2019|H5N2|2019-11-04
4071029|A/Mallard(Anas_platyrhynchos)/South_Korea/KNU2019-20/2019|H5N1|2019-03-03
389829|A/Avian/South_Korea/K58/2018|H5N3|2018-08-21
819124|A/mallard/Netherlands/19001282-001/2019|H5N1|2019-01-23
18537218|A/Mallard/Netherlands/96/2019|H5N3|2019-12-09
1263541|A/mallard/France/20P017917/2020|H5N3|2020-12-07
11402318|A/turkey/England/018179/2021|H5N3|2021-03-25
12853041|A/ostrich/South_Africa/090581/2020|H5N2|2020-09-18
822837|A/green_sandpiper/Netherlands/19011490-013/2019|H5N1|2019-08-13
17978925|A/mallard/Poland/P077w1/2018|H5N2|2018-09-12
2234808|A/knot_wader/Ireland/473_21VIR2956-2/2021|H5N3|2021-01-01
2234807|A/knot_wader/Ireland/472_21VIR2956-1/2021|H5N3|2021-01-01
1259399|A/red_knot/France/21P003249/2021|H5N3|2021-01-18
1259397|A/curlew/France/21P003648/2021|H5N3|2021-01-22
1063993|A/common_kestrel/Denmark/16023-01/2021-1.01(H5N3)|H5N3|2021-01-01
2172530|A/common_buzzard/Netherlands/21024357-002/2021|H5N3|2021-02-22
1206285|A/red_knot/Germany-NI/AI00394/2020|H5N3|2020-12-30
1205487|A/red_knot/Germany-SH/AI03421/2020|H5N3|2020-12-14
1205485|A/red_knot/Germany-SH/AI03419/2020|H5N3|2020-12-14
11392591|A/peregrine_falcon/Northern_Ireland/AI102021-2/2021|H5N3|2021-01-22
2234831|A/peregrine_falcon/Northern_Ireland/21VIR1662-3/2021|H5N3|2021-01-25
1048239|A/common_buzzard/Netherlands/21021023-002/2021|H5N3|2021-01-04
1205489|A/red_knot/Germany-SH/AI03424/2020|H5N3|2020-12-16
661312|A/guinea_fowl/Germany-NW/AI01184/2020|H5N8|2020-09-07
9261740|A/Mallard/Netherlands/17/2021|H5N3|2021-12-29
14768234|A/wild_waterbird/Queensland/JCU-272/2019|H5N9|2019-08-09
A/turkey/Germany-BB/AI00868/2021|mixH5N8-3|2021-01-22
A/turkey/Germany-BB/AI00868/2021|mixH5N8-2|2021-01-22
502593|A/Anas_Platyrhynchos/Belgium/10811_6/2019|H5N6|2019-09-21
13243697|A/environment/England/030642/2020|H5N2|2020-10-31
3031133|A_mallard_Denmark_12946-11_2020-10-26|H5N2|2020-10-26
5863749|A/green-winged-teal/Georgia/DT-22894/2020|H5N3|2020-10-20
A/turkey/Germany-BB/AI00868/2021|mixH5N8-4|2021-01-22
7595492|A/Anas_platyrhynchos/Belgium/10413_0003/2020|H5N2|2020-10-04
3161955|A/whooper_swan/Henan/T578/2019|H5N2|2019-01-15
3161847|A/whooper_swan/Henan/T258/2019|H5N2|2019-01-15
403722|A/Spot-billed_Duck/Russia_Primorje/226/2019|H5N3|2019-10-20
4063594|A/duck/Tottori/Vac-T1/2020|H5N6|2020-05-28
14769870|A/wild_waterbird/Western_Australia/AS19-1358-170/2019|H5N3|2019-04-02
14769871|A/wild_waterbird/Western_Australia/AS19-1358-178/2019|H5N7|2019-04-02
14870959|A/chicken/Queretaro/CPA-04673-1/2019|H5N2|2019-03-20
393518|A/Avian/Iran/38AMA/2019|H5N3|2019-03-30
0.06

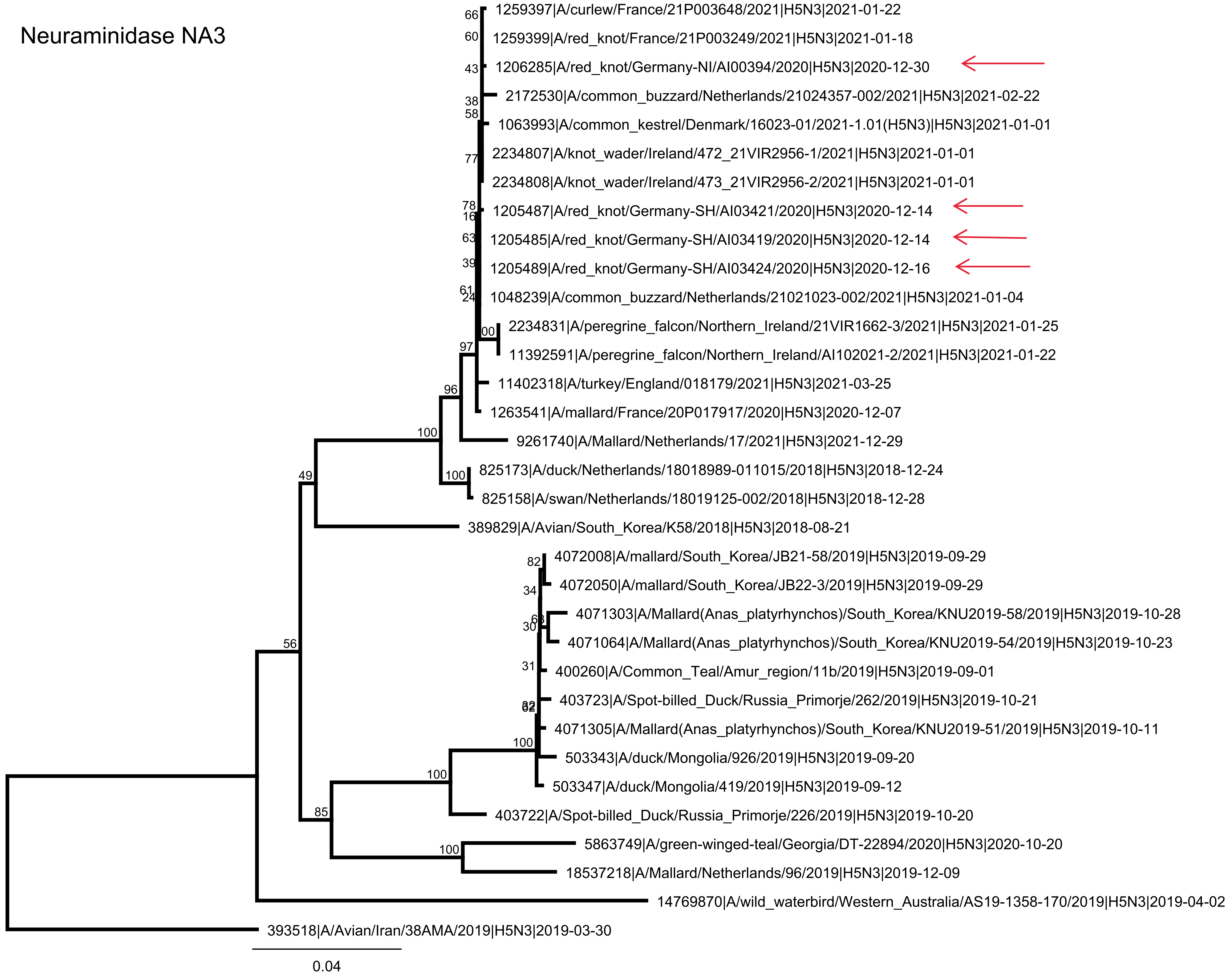
Neuraminidase NA3
1259397|A/curlew/France/21P003648/2021|H5N3|2021-01-22
1259399|A/red_knot/France/21P003249/2021|H5N3|2021-01-18
1206285|A/red_knot/Germany-NI/AI00394/2020|H5N3|2020-12-30
2172530|A/common_buzzard/Netherlands/21024357-002/2021|H5N3|2021-02-22
1063993|A/common_kestrel/Denmark/16023-01/2021-1.01(H5N3)|H5N3|2021-01-01
2234807|A/knot_wader/Ireland/472_21VIR2956-1/2021|H5N3|2021-01-01
2234808|A/knot_wader/Ireland/473_21VIR2956-2/2021|H5N3|2021-01-01
1205487|A/red_knot/Germany-SH/AI03421/2020|H5N3|2020-12-14
1205485|A/red_knot/Germany-SH/AI03419/2020|H5N3|2020-12-14
1205489|A/red_knot/Germany-SH/AI03424/2020|H5N3|2020-12-16
1048239|A/common_buzzard/Netherlands/21021023-002/2021|H5N3|2021-01-04
2234831|A/peregrine_falcon/Northern_Ireland/21VIR1662-3/2021|H5N3|2021-01-25
11392591|A/peregrine_falcon/Northern_Ireland/AI102021-2/2021|H5N3|2021-01-22
11402318|A/turkey/England/018179/2021|H5N3|2021-03-25
1263541|A/mallard/France/20P017917/2020|H5N3|2020-12-07
9261740|A/Mallard/Netherlands/17/2021|H5N3|2021-12-29
825173|A/duck/Netherlands/18018989-011015/2018|H5N3|2018-12-24
825158|A/swan/Netherlands/18019125-002/2018|H5N3|2018-12-28
389829|A/Avian/South_Korea/K58/2018|H5N3|2018-08-21
4072008|A/mallard/South_Korea/JB21-58/2019|H5N3|2019-09-29
4072050|A/mallard/South_Korea/JB22-3/2019|H5N3|2019-09-29
4071303|A/Mallard(Anas_platyrhynchos)/South_Korea/KNU2019-58/2019|H5N3|2019-10-28
4071064|A/Mallard(Anas_platyrhynchos)/South_Korea/KNU2019-54/2019|H5N3|2019-10-23
400260|A/Common_Teal/Amur_region/11b/2019|H5N3|2019-09-01
403723|A/Spot-billed_Duck/Russia_Primorje/262/2019|H5N3|2019-10-21
4071305|A/Mallard(Anas_platyrhynchos)/South_Korea/KNU2019-51/2019|H5N3|2019-10-11
503343|A/duck/Mongolia/926/2019|H5N3|2019-09-20
503347|A/duck/Mongolia/419/2019|H5N3|2019-09-12
403722|A/Spot-billed_Duck/Russia_Primorje/226/2019|H5N3|2019-10-20
5863749|A/green-winged-teal/Georgia/DT-22894/2020|H5N3|2020-10-20
18537218|A/Mallard/Netherlands/96/2019|H5N3|2019-12-09
14769870|A/wild_waterbird/Western_Australia/AS19-1358-170/2019|H5N3|2019-04-02
393518|A/Avian/Iran/38AMA/2019|H5N3|2019-03-30
0.04

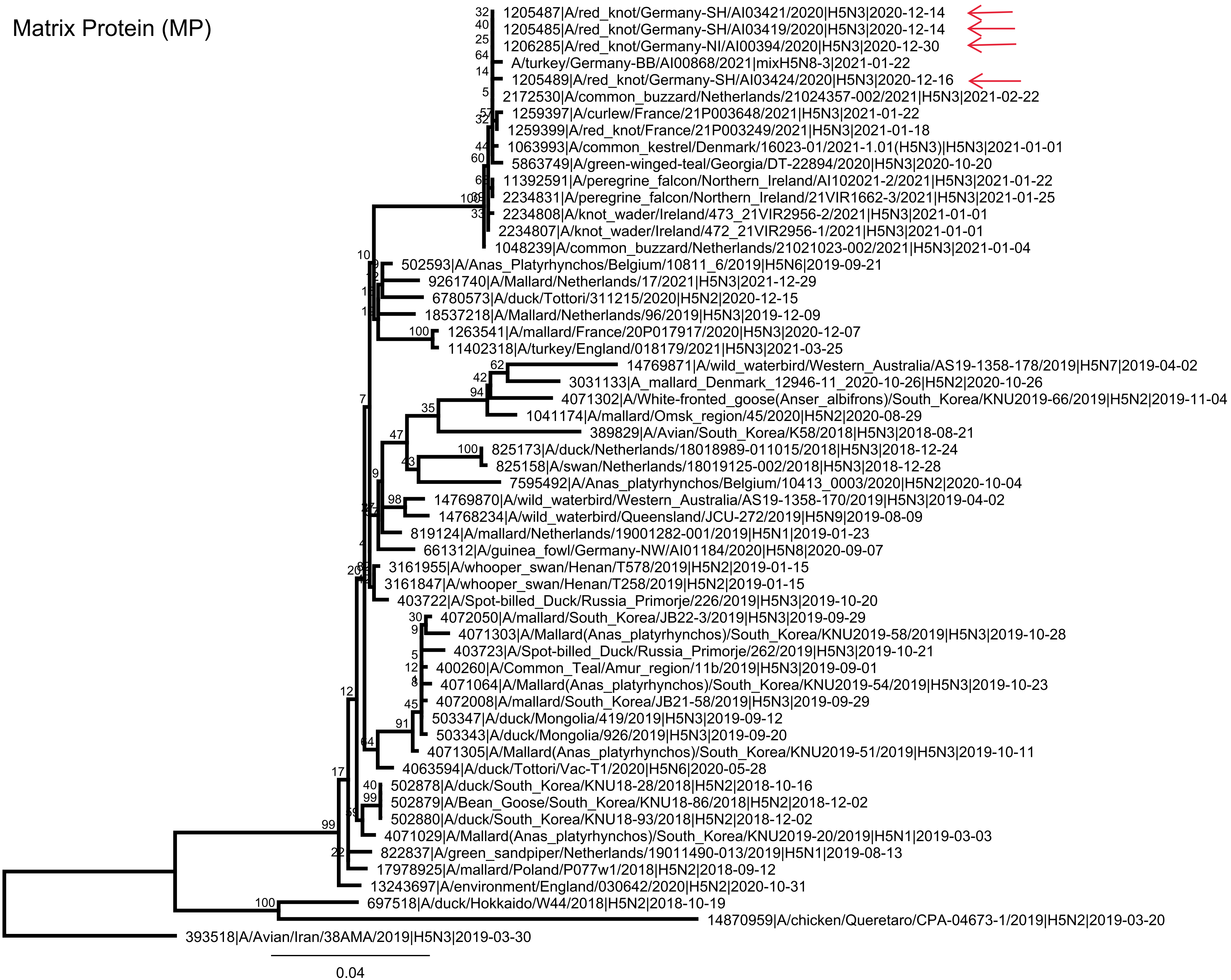
Matrix Protein (MP)
1205487|A/red_knot/Germany-SH/AI03421/2020|H5N3|2020-12-14
1205485|A/red_knot/Germany-SH/AI03419/2020|H5N3|2020-12-14
1206285|A/red_knot/Germany-NI/AI00394/2020|H5N3|2020-12-30
A/turkey/Germany-BB/AI00868/2021|mixH5N8-3|2021-01-22
1205489|A/red_knot/Germany-SH/AI03424/2020|H5N3|2020-12-16
2172530|A/common_buzzard/Netherlands/21024357-002/2021|H5N3|2021-02-22
1259397|A/curlew/France/21P003648/2021|H5N3|2021-01-22
1259399|A/red_knot/France/21P003249/2021|H5N3|2021-01-18
1063993|A/common_kestrel/Denmark/16023-01/2021-1.01(H5N3)|H5N3|2021-01-01
5863749|A/green-winged-teal/Georgia/DT-22894/2020|H5N3|2020-10-20
11392591|A/peregrine_falcon/Northern_Ireland/AI102021-2/2021|H5N3|2021-01-22
2234831|A/peregrine_falcon/Northern_Ireland/21VIR1662-3/2021|H5N3|2021-01-25
2234808|A/knot_wader/Ireland/473_21VIR2956-2/2021|H5N3|2021-01-01
2234807|A/knot_wader/Ireland/472_21VIR2956-1/2021|H5N3|2021-01-01
1048239|A/common_buzzard/Netherlands/21021023-002/2021|H5N3|2021-01-04
502593|A/Anas_Platyrhynchos/Belgium/10811_6/2019|H5N6|2019-09-21
9261740|A/Mallard/Netherlands/17/2021|H5N3|2021-12-29
6780573|A/duck/Tottori/311215/2020|H5N2|2020-12-15
18537218|A/Mallard/Netherlands/96/2019|H5N3|2019-12-09
1263541|A/mallard/France/20P017917/2020|H5N3|2020-12-07
11402318|A/turkey/England/018179/2021|H5N3|2021-03-25
14769871|A/wild_waterbird/Western_Australia/AS19-1358-178/2019|H5N7|2019-04-02
3031133|A_mallard_Denmark_12946-11_2020-10-26|H5N2|2020-10-26
4071302|A/White-fronted_goose(Anser_albifrons)/South_Korea/KNU2019-66/2019|H5N2|2019-11-04
1041174|A/mallard/Omsk_region/45/2020|H5N2|2020-08-29
389829|A/Avian/South_Korea/K58/2018|H5N3|2018-08-21
825173|A/duck/Netherlands/18018989-011015/2018|H5N3|2018-12-24
825158|A/swan/Netherlands/18019125-002/2018|H5N3|2018-12-28
7595492|A/Anas_platyrhynchos/Belgium/10413_0003/2020|H5N2|2020-10-04
14769870|A/wild_waterbird/Western_Australia/AS19-1358-170/2019|H5N3|2019-04-02
14768234|A/wild_waterbird/Queensland/JCU-272/2019|H5N9|2019-08-09
819124|A/mallard/Netherlands/19001282-001/2019|H5N1|2019-01-23
661312|A/guinea_fowl/Germany-NW/AI01184/2020|H5N8|2020-09-07
3161955|A/whooper_swan/Henan/T578/2019|H5N2|2019-01-15
3161847|A/whooper_swan/Henan/T258/2019|H5N2|2019-01-15
403722|A/Spot-billed_Duck/Russia_Primorje/226/2019|H5N3|2019-10-20
4072050|A/mallard/South_Korea/JB22-3/2019|H5N3|2019-09-29
4071303|A/Mallard(Anas_platyrhynchos)/South_Korea/KNU2019-58/2019|H5N3|2019-10-28
403723|A/Spot-billed_Duck/Russia_Primorje/262/2019|H5N3|2019-10-21
400260|A/Common_Teal/Amur_region/11b/2019|H5N3|2019-09-01
4071064|A/Mallard(Anas_platyrhynchos)/South_Korea/KNU2019-54/2019|H5N3|2019-10-23
4072008|A/mallard/South_Korea/JB21-58/2019|H5N3|2019-09-29
503347|A/duck/Mongolia/419/2019|H5N3|2019-09-12
503343|A/duck/Mongolia/926/2019|H5N3|2019-09-20
4071305|A/Mallard(Anas_platyrhynchos)/South_Korea/KNU2019-51/2019|H5N3|2019-10-11
4063594|A/duck/Tottori/Vac-T1/2020|H5N6|2020-05-28
502878|A/duck/South_Korea/KNU18-28/2018|H5N2|2018-10-16
502879|A/Bean_Goose/South_Korea/KNU18-86/2018|H5N2|2018-12-02
502880|A/duck/South_Korea/KNU18-93/2018|H5N2|2018-12-02
4071029|A/Mallard(Anas_platyrhynchos)/South_Korea/KNU2019-20/2019|H5N1|2019-03-03
822837|A/green_sandpiper/Netherlands/19011490-013/2019|H5N1|2019-08-13
17978925|A/mallard/Poland/P077w1/2018|H5N2|2018-09-12
13243697|A/environment/England/030642/2020|H5N2|2020-10-31
697518|A/duck/Hokkaido/W44/2018|H5N2|2018-10-19
14870959|A/chicken/Queretaro/CPA-04673-1/2019|H5N2|2019-03-20
393518|A/Avian/Iran/38AMA/2019|H5N3|2019-03-30
0.04

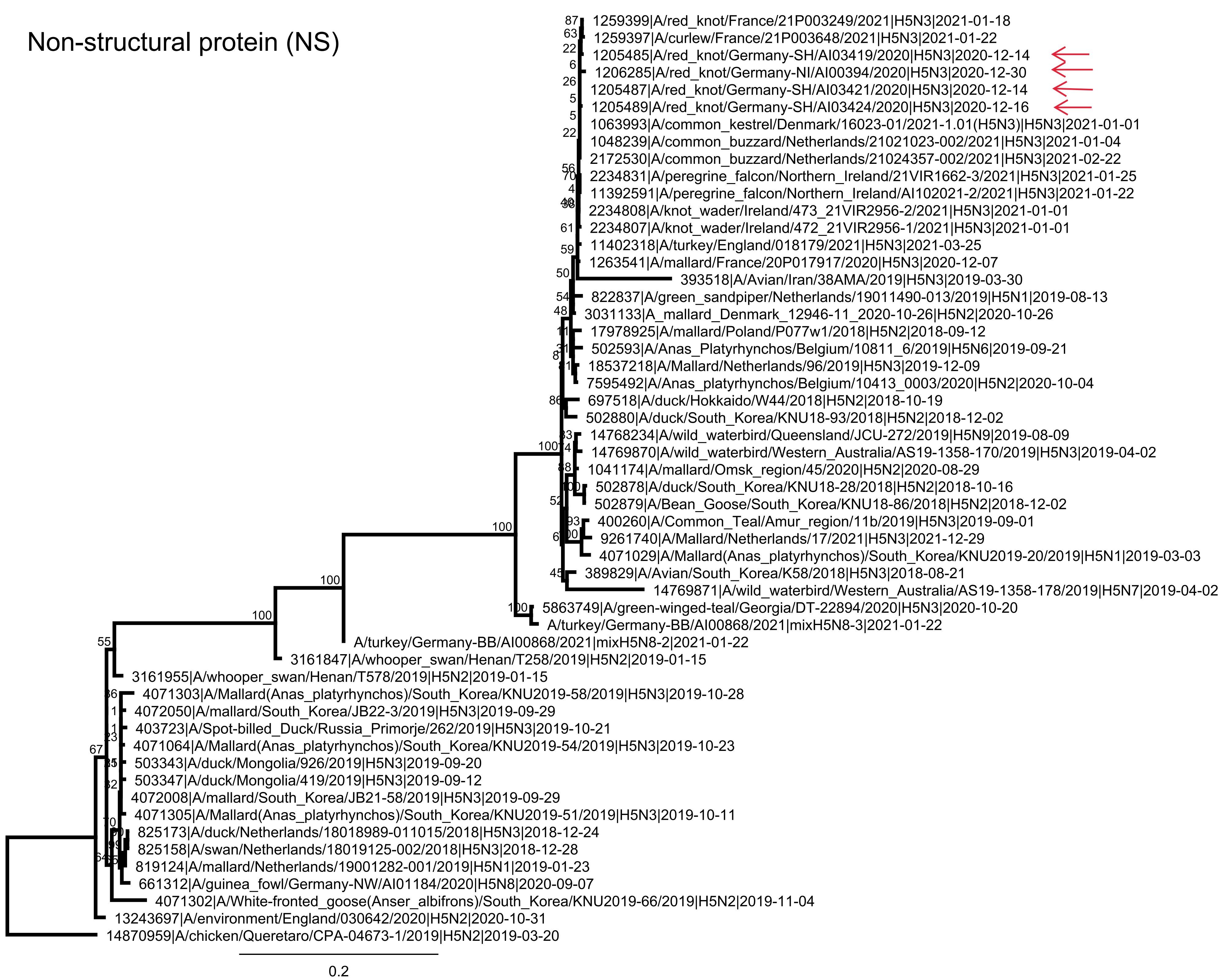
Non-structural protein (NS)
1259399|A/red_knot/France/21P003249/2021|H5N3|2021-01-18
1259397|A/curlew/France/21P003648/2021|H5N3|2021-01-22
1205485|A/red_knot/Germany-SH/AI03419/2020|H5N3|2020-12-14
1206285|A/red_knot/Germany-NI/AI00394/2020|H5N3|2020-12-30
1205487|A/red_knot/Germany-SH/AI03421/2020|H5N3|2020-12-14
1205489|A/red_knot/Germany-SH/AI03424/2020|H5N3|2020-12-16
1063993|A/common_kestrel/Denmark/16023-01/2021-1.01(H5N3)|H5N3|2021-01-01
1048239|A/common_buzzard/Netherlands/21021023-002/2021|H5N3|2021-01-04
2172530|A/common_buzzard/Netherlands/21024357-002/2021|H5N3|2021-02-22
2234831|A/peregrine_falcon/Northern_Ireland/21VIR1662-3/2021|H5N3|2021-01-25
11392591|A/peregrine_falcon/Northern_Ireland/AI102021-2/2021|H5N3|2021-01-22
2234808|A/knot_wader/Ireland/473_21VIR2956-2/2021|H5N3|2021-01-01
2234807|A/knot_wader/Ireland/472_21VIR2956-1/2021|H5N3|2021-01-01
11402318|A/turkey/England/018179/2021|H5N3|2021-03-25
1263541|A/mallard/France/20P017917/2020|H5N3|2020-12-07
393518|A/Avian/Iran/38AMA/2019|H5N3|2019-03-30
822837|A/green_sandpiper/Netherlands/19011490-013/2019|H5N1|2019-08-13
3031133|A_mallard_Denmark_12946-11_2020-10-26|H5N2|2020-10-26
17978925|A/mallard/Poland/P077w1/2018|H5N2|2018-09-12
502593|A/Anas_Platyrhynchos/Belgium/10811_6/2019|H5N6|2019-09-21
18537218|A/Mallard/Netherlands/96/2019|H5N3|2019-12-09
7595492|A/Anas_platyrhynchos/Belgium/10413_0003/2020|H5N2|2020-10-04
697518|A/duck/Hokkaido/W44/2018|H5N2|2018-10-19
502880|A/duck/South_Korea/KNU18-93/2018|H5N2|2018-12-02
14768234|A/wild_waterbird/Queensland/JCU-272/2019|H5N9|2019-08-09
14769870|A/wild_waterbird/Western_Australia/AS19-1358-170/2019|H5N3|2019-04-02
1041174|A/mallard/Omsk_region/45/2020|H5N2|2020-08-29
502878|A/duck/South_Korea/KNU18-28/2018|H5N2|2018-10-16
502879|A/Bean_Goose/South_Korea/KNU18-86/2018|H5N2|2018-12-02
400260|A/Common_Teal/Amur_region/11b/2019|H5N3|2019-09-01
9261740|A/Mallard/Netherlands/17/2021|H5N3|2021-12-29
4071029|A/Mallard(Anas_platyrhynchos)/South_Korea/KNU2019-20/2019|H5N1|2019-03-03
389829|A/Avian/South_Korea/K58/2018|H5N3|2018-08-21
14769871|A/wild_waterbird/Western_Australia/AS19-1358-178/2019|H5N7|2019-04-02
5863749|A/green-winged-teal/Georgia/DT-22894/2020|H5N3|2020-10-20
A/turkey/Germany-BB/AI00868/2021|mixH5N8-3|2021-01-22
A/turkey/Germany-BB/AI00868/2021|mixH5N8-2|2021-01-22
3161847|A/whooper_swan/Henan/T258/2019|H5N2|2019-01-15
3161955|A/whooper_swan/Henan/T578/2019|H5N2|2019-01-15
4071303|A/Mallard(Anas_platyrhynchos)/South_Korea/KNU2019-58/2019|H5N3|2019-10-28
4072050|A/mallard/South_Korea/JB22-3/2019|H5N3|2019-09-29
403723|A/Spot-billed_Duck/Russia_Primorje/262/2019|H5N3|2019-10-21
4071064|A/Mallard(Anas_platyrhynchos)/South_Korea/KNU2019-54/2019|H5N3|2019-10-23
503343|A/duck/Mongolia/926/2019|H5N3|2019-09-20
503347|A/duck/Mongolia/419/2019|H5N3|2019-09-12
4072008|A/mallard/South_Korea/JB21-58/2019|H5N3|2019-09-29
4071305|A/Mallard(Anas_platyrhynchos)/South_Korea/KNU2019-51/2019|H5N3|2019-10-11
825173|A/duck/Netherlands/18018989-011015/2018|H5N3|2018-12-24
825158|A/swan/Netherlands/18019125-002/2018|H5N3|2018-12-28
819124|A/mallard/Netherlands/19001282-001/2019|H5N1|2019-01-23
661312|A/guinea_fowl/Germany-NW/AI01184/2020|H5N8|2020-09-07
4071302|A/White-fronted_goose(Anser_albifrons)/South_Korea/KNU2019-66/2019|H5N2|2019-11-04
13243697|A/environment/England/030642/2020|H5N2|2020-10-31
14870959|A/chicken/Queretaro/CPA-04673-1/2019|H5N2|2019-03-20
0.2

***Data source: sequence information with virus name, accession and metadata. We acknowledge the listed originating and submitting laboratories for sharing their data.***

| virus name | accession | Subtype | Submitting Lab | Originating Lab |
|---|---|---|---|---|
| A/Avian/South Korea/K58/2018 | 389829 | H5N3 | | |
| A/Avian/Iran/38AMA/2019 | 393518 | H5N3 | | |
| A/Common Teal/Amur region/11b/2019 | 400260 | H5N3 | National Institute of Animal Health | Research Institute of Experimental and Clinical Medicine |
| A/Spot-billed Duck/Russia Primorje/226/2019 | 403722 | H5N3 | National Institute of Animal Health | Research Institute of Experimental and Clinical Medicine |
| A/Spot-billed Duck/Russia Primorje/262/2019 | 403723 | H5N3 | National Institute of Animal Health | Research Institute of Experimental and Clinical Medicine |
| A/chicken/Denmark/S02750-3/2020 | 410987 | H5N1 | Statens Serum Institute | Statens Serumm Institute |
| A/Anas Platyrhynchos/Belgium/10811 6/2019 | 502593 | H5N6 | | |
| A/duck/South Korea/KNU18-28/2018 | 502878 | H5N2 | | |
| A/Bean Goose/South Korea/KNU18-86/2018 | 502879 | H5N2 | | |
| A/duck/South Korea/KNU18-93/2018 | 502880 | H5N2 | | |
| A/duck/Mongolia/926/2019 | 503343 | H5N3 | | |
| A/duck/Mongolia/419/2019 | 503347 | H5N3 | | |
| A/guinea fowl/Germany-NW/AI01184/2020 | 661312 | H5N8 | Friedrich-Loeffler-Institut | Chemisches und Veterinäruntersuchungsamt Münsterland-Emscher-Lippe |
| A/duck/Hokkaido/W44/2018 | 697518 | H5N2 | | |
| A/mallard/Netherlands/19001282-001/2019 | 819124 | H5N1 | Wageningen Bioveterinary Research | Wageningen Bioveterinary Research |
| A/green sandpiper/Netherlands/19011490-013/2019 | 822837 | H5N1 | Wageningen Bioveterinary Research | Wageningen Bioveterinary Research |
| A/swan/Netherlands/18019125-002/2018 | 825158 | H5N3 | Wageningen Bioveterinary Research | Wageningen Bioveterinary Research |
| A/duck/Netherlands/18018989-011015/2018 | 825173 | H5N3 | Wageningen Bioveterinary Research | Wageningen Bioveterinary Research |
| A/mallard/Omsk region/45/2020 | 1041174 | H5N2 | National Institute of Animal Health | Research Institute of Experimental and Clinical Medicine |
| A/common buzzard/Netherlands/21021023-002/2021 | 1048239 | H5N3 | Wageningen Bioveterinary Research | Wageningen Bioveterinary Research |
| A/common kestrel/Denmark/16023-01/2021-01-01 | 1063993 | H5N3 | Statens Serum Institute | Statens Serumm Institute |
| A/common kestrel/Denmark/16023-01/2021-1.01(H5N3) | 1063993 | H5N3 | Statens Serum Institute | Statens Serumm Institute |
| A/red knot/Germany-SH/AI03419/2020 | 1205485 | H5N3 | Friedrich-Loeffler-Institut | Landeslabor Schleswig-Holstein |
| A/red knot/Germany-SH/AI03421/2020 | 1205487 | H5N3 | Friedrich-Loeffler-Institut | Landeslabor Schleswig-Holstein |
| A/red knot/Germany-SH/AI03424/2020 | 1205489 | H5N3 | Friedrich-Loeffler-Institut | Landeslabor Schleswig-Holstein |
| A/red knot/Germany-NI/AI00394/2020 | 1206285 | H5N3 | Friedrich-Loeffler-Institut | Lebensmittel- und Veterinärinstitut Oldenburg - Standort Veterinärinstitut |
| A/curlew/France/21P003648/2021 | 1259397 | H5N3 | ANSES Agence Nationale De Securite Sanitaire De L'alimentation | Anses (Ploufragan-Plouzané) |
| A/red knot/France/21P003249/2021 | 1259399 | H5N3 | ANSES Agence Nationale De Securite Sanitaire De L'alimentation | Anses (Ploufragan-Plouzané) |
| A/mallard/France/20P017917/2020 | 1263541 | H5N3 | ANSES Agence Nationale De Securite Sanitaire De L'alimentation | Anses (Ploufragan-Plouzané) |
| A/common buzzard/Netherlands/21024357-002/2021 | 2172530 | H5N3 | Wageningen Bioveterinary Research | Wageningen Bioveterinary Research |
| A/knot wader/Ireland/472 21VIR2956-1/2021 | 2234807 | H5N3 | Istituto Zooprofilattico Sperimentale Delle Venezie | Istituto Zooprofilattico Sperimentale delle Venezie, EU/OIE/Reference Laboratory and FAO Reference Centre for AI and ND |
| A/knot wader/Ireland/473 21VIR2956-2/2021 | 2234808 | H5N3 | Istituto Zooprofilattico Sperimentale Delle Venezie | Istituto Zooprofilattico Sperimentale delle Venezie, EU/OIE/Reference Laboratory and FAO Reference Centre for AI and ND |
| A/peregrine falcon/Northern Ireland/21VIR1662-3/2021 | 2234831 | H5N3 | Istituto Zooprofilattico Sperimentale Delle Venezie | Istituto Zooprofilattico Sperimentale delle Venezie, EU/OIE/Reference Laboratory and FAO Reference Centre for AI and ND |
| A mallard Denmark 12946-11 2020-10-26 | 3031133 | H5N2 | Statens Serum Institute | Statens Serumm Institute |
| A/whooper swan/Henan/T258/2019 | 3161847 | H5N2 | Chinese Academy of Forestry | Xi'an Tianlong Science and Technology Co., Ltd. |
| A/whooper swan/Henan/T578/2019 | 3161955 | H5N2 | Chinese Academy of Forestry | Xi'an Tianlong Science and Technology Co., Ltd. |
| A/duck/Tottori/Vac-T1/2020 | 4063594 | H5N6 | | |
| A/Mallard(Anas platyrhynchos)/South Korea/KNU2019-20/2019 | 4071029 | H5N1 | | |
| A/Mallard(Anas platyrhynchos)/South Korea/KNU2019-54/2019 | 4071064 | H5N3 | | |
| A/White-fronted goose(Anser albifrons)/South Korea/KNU2019-66/2019 | 4071302 | H5N2 | | |
| A/Mallard(Anas platyrhynchos)/South Korea/KNU2019-58/2019 | 4071303 | H5N3 | | |
| A/Mallard(Anas platyrhynchos)/South Korea/KNU2019-51/2019 | 4071305 | H5N3 | | |
| A/mallard/South Korea/JB21-58/2019 | 4072008 | H5N3 | | |
| A/mallard/South Korea/JB22-3/2019 | 4072050 | H5N3 | | |
| A/turkey/Germany-BB/AI00868/2021 | 5095644 | H5N8 | Friedrich-Loeffler-Institut | Landeslabor Berlin-Brandenburg |
| A/green-winged-teal/Georgia/DT-22894/2020 | 5863749 | H5N3 | Royal Veterinary College | Erasmus Medical Center |
| A/duck/Tottori/311215/2020 | 6780573 | H5N2 | | |
| A/Anas platyrhynchos/Belgium/10413 0003/2020 | 7595492 | H5N2 | Sciensano, Department of Animal Infectious Diseases | Sciensano - Animal Infectious Diseases |
| A/Mallard/Netherlands/17/2021 | 9261740 | H5N3 | Erasmus Medical Center | Erasmus Medical Center |
| A/peregrine falcon/Northern Ireland/AI102021-2/2021 | 11392591 | H5N3 | Animal and Plant Health Agency (APHA) | Animal and Plant Health Agency (APHA) |
| A/turkey/England/018179/2021 | 11402318 | H5N3 | Animal and Plant Health Agency (APHA) | Animal and Plant Health Agency (APHA) |
| A/ostrich/South Africa/090581/2020 | 12853041 | H5N2 | University of Pretoria | Western Cape Provincial Veterinary Laboratory |
| A/environment/England/030642/2020 | 13243697 | H5N2 | Animal and Plant Health Agency (APHA) | Animal and Plant Health Agency (APHA) |
| A/wild waterbird/Queensland/JCU-272/2019 | 14768234 | H5N9 | | |
| A/wild waterbird/Western Australia/AS19-1358-170/2019 | 14769870 | H5N3 | | |
| A/wild waterbird/Western Australia/AS19-1358-178/2019 | 14769871 | H5N7 | | |
| A/Northern Pintail/Tottori/NIES1378/2020 | 14867043 | H5N2 | | |
| A/chicken/Queretaro/CPA-04673-1/2019 | 14870959 | H5N2 | | |
| A/duck/Vietnam/HN5074/2018 | 17734171 | H5N3 | | |
| A/duck/Vietnam/HN5076/2018 | 17734177 | H5N3 | | |
| A/duck/Vietnam/HN5074/2018 | 17768804 | H5N3 | | Center for Research on Influenza Pathogenesis |
| A/duck/Vietnam/HN5076/2018 | 17768805 | H5N3 | | Center for Research on Influenza Pathogenesis |
| A/mallard/Poland/P077w1/2018 | 17978925 | H5N2 | National Veterinary Research Institute | National Veterinary Research Institut Poland, PIWet-PIB |
| A/Mallard/Netherlands/96/2019 | 18537218 | H5N3 | Erasmus Medical Center | Erasmus Medical Center |
